# Molecular cascades and cell-type specific signatures in ASD revealed by single cell genomics

**DOI:** 10.1101/2023.03.10.530869

**Authors:** Brie Wamsley, Lucy Bicks, Yuyan Cheng, Riki Kawaguchi, Diana Quintero, Jennifer Grundman, Jianyin Liu, Shaohua Xiao, Natalie Hawken, Michael Margolis, Samantha Mazariegos, Daniel H. Geschwind

## Abstract

Understanding how genetic variation exerts its effects on the human brain in health and disease has been greatly informed by functional genomic characterization. Studies over the last decade have demonstrated robust evidence of convergent transcriptional and epigenetic profiles in post-mortem cerebral cortex from individuals with Autism Spectrum Disorder (ASD). Here, we perform deep single nuclear (sn) RNAseq to elucidate changes in cell composition, cellular transcriptomes and putative candidate drivers associated with ASD, which we corroborate using snATAC-seq and spatial profiling. We find changes in cell state composition representing transitions from homeostatic to reactive profiles in microglia and astrocytes, a pattern extending to oligodendrocytes and blood brain barrier cells. We identify profound changes in differential expression involving thousands of genes across neuronal and glial subtypes, of which a substantial portion can be accounted for by specific transcription factor networks that are significantly enriched in common and rare genetic risk for ASD. These data, which are available as part of the PsychENCODE consortium, provide robust causal anchors and resultant molecular phenotypes for understanding ASD changes in human brain.

**One-Sentence Summary:** We define the molecular cascades and cells disrupted in post-mortem brain in ASD by performing spatial, single nuclear RNA, and epigenetic profiling, and characterize, at unmatched resolution, the functional regulation of cell-type specific signatures underlying the molecular differences and physiology of ASD.

**Main Text:** Psychiatric disorders are defined primarily by behavioral and cognitive characteristics and are classically distinguished from neurological disorders by lacking the associated visible histological or macroscopic pathology observed in neurological conditions. However, a growing body of evidence based on genomic profiling reveals consistent molecular differences in brain tissue from specific neuropsychiatric conditions compared with brain tissue from neurotypical individuals (1–5). In Autism Spectrum Disorder (ASD) robust transcriptomic and epigenetic alterations in the cerebral cortex from patients have been documented over the last decade, delineating a reproducible pattern of molecular pathology (5–13). This robust molecular signature obtained primarily from transcriptomic profiling of bulk cortical tissue has identified convergent biological pathways in ASD brain, which is characterized by an upregulation of immune signaling genes, downregulation of specific neuronal markers, synaptic genes, and an attenuation of the typical patterns of gene expression associated with cortical regional identity (6–8,12–14).

These genomic data represent an essential lens through which to understand the cellular and physiological changes occurring in the brains of autistic individuals and to describe potential causal mechanisms via their integration with genetic risk variants (1,4,5). However, small sample sizes in the case of single cell analysis (13), or profiling restricted to bulk tissue have limited biological insights as to the differences in laminar, circuit level, and cell-type specific pathways affected in ASD, as well as their underlying gene regulatory mechanisms. To address these limitations, we leveraged improvement in single cell analyses to profile the largest ASD cohort to date, consisting of 64 cases and controls. This resource, generated as a core component of the PsychENCODE consortium (1,2,4,5,8; http://www.psychencode.org), also enables us to characterize underlying candidate regulatory mechanisms and to connect causal drivers with the observed changes at a cell-type specific level in ASD, providing a deeper and more generalizable understanding of the cell types and biological mechanisms that underlie ASD. These data are available via PsychEncode portals for download (http://psychencode.synapse.org) and on the PsychSCREEN browser (*in development*, http://psychscreen.beta.wenglab.org).

## Results

### Single cell genomic analysis of human ASD and control cortex

We generated snRNAseq from the human frontal cortex (PFC, Brodmann areas 9 and 4/6) from 33 ASD and 31 control subjects (Fig. 1A; Fig. S1A; Table S1-1), profiling on average more than 10,000 cells per individual (Fig. S1A,C). We include 5 individuals with a monogenic form of ASD, 15q dup syndrome (Table S1-1), which we had previously shown to mirror transcriptomic changes observed in bulk tissue from cases of idiopathic ASD (7,8). Individual samples ranged from ages 2 to 60 (mean age: 20) and were selected to be matched by age, sex, and cause of death balanced across cases and controls (Fig. S1B,C; Table S1-1). We isolated nuclei and prepped cells for snRNAseq using the 10x genomics platform (Methods), generating over 800,000 nuclei, making it the largest sample by ≈ 8-fold (Table S1-4). Following quality control and doublet removal, we analyzed 591,043 nuclei transcriptional profiles representing 306,347 cells from ASD and 284,696 cells from control subjects (average read depth of 6.6 × 10^4^ per cell) with a mean of 1860 genes and 4164 RNA molecules detected per cell (Fig. S1C; Table S2). Additionally, we simultaneously isolated nuclei to generate chromatin conformation states via single-nuclei ATAC-seq in 22 cases and controls to validate predicted cell-type specific regulatory networks via identification of enriched transcription factor (TF) binding sites and TF foot-printing detailed further below (Fig. 1A).

**Figure 1.**
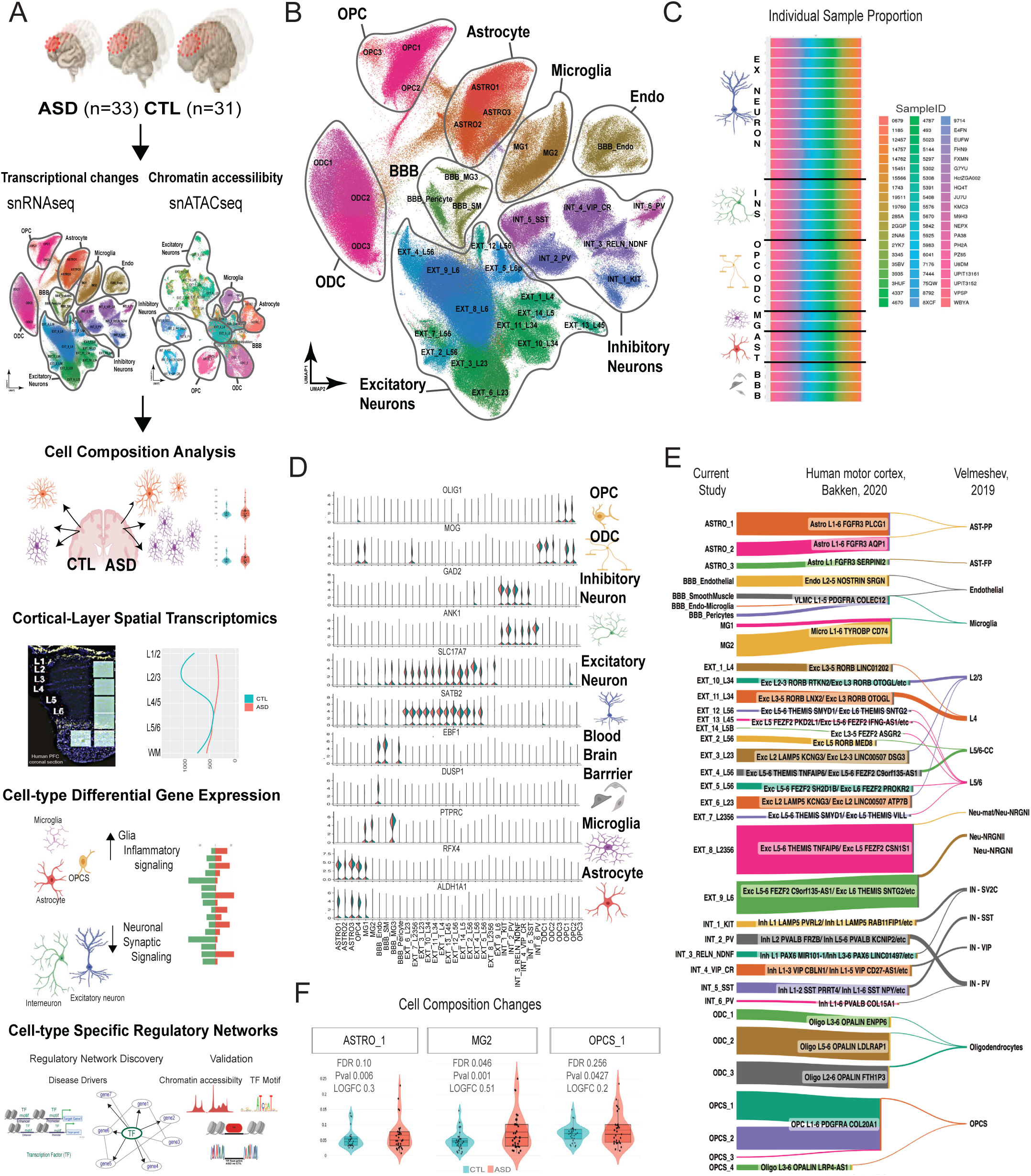
Single-cell analysis of the ASD cortex. A) Schematic of the major components of the study: the frontal cortex (BA9/4/6) of the largest cohort of ASD and CTL individuals were subjected to snRNAseq - 1/3 of these samples were also subjected to snATACseq. Samples were then analyzed for changes in cell composition, and spatially resolved transcriptomics was conducted to assess laminar level changes in cell-types and differential gene expression. We generate regulatory networks (SCENIC) discovered to drive cell-type specific changes in DE and validate these changes with TF foot printing (TOBIAS) within open chromatin changes (snATACseq). B) UMAP of cell clusters defined by snRNAseq C) Individual sample proportions of cell clusters shows even distribution across clusters. D) Stacked violin plots of marker gene expression in all clusters for *OLIG1* for OPCs, MOG for ODCs (and OPCS), *GAD2* and *ANK1* for INs, *SLC17A7* and *SATB2* for Ex neurons, *EBF1* and *DUSP1* to mark pericytes, smooth muscle, and endothelial cells of the BBB, *PTPRC* for microglia, *RFX4* and *ALDH1A1* for Astrocytes. Violin plots are split colored to mark CTL expression in blue and ASD expression in red. E) Sankey plot of cell cluster equivalents from this current study, left, to reference cell types from Allen brain atlas (19), human motor cortex, center, and Velmeshev et al. (13), human prefrontal cortex, right. The height of the colored bar for each cell type depicts the number of cells from this current study from the left to center; the height of bar from the center to the right represents the number of cells from Velmeshev (13) cluster types. The cluster names from Allen atlas (19) were shortened to the top matching clusters - see Table S1-3 for all cluster name assignments. F) Significant increase in reactive astrocytes and microglia in cortex from individuals with ASD. Cell proportions for ASTRO1 (*FDR* = *0.10 P = 0.006, LOGFC = 0.3*), left, MG2 (*FDR* = *0.046 P = 0.001, LOGFC = 0.51*), center, and OPCS_1. (*FDR = 0.26, P. = 0.043, LOGFC = 0.2*), right from CTL individuals (blue) and ASD individuals (red). *Abbreviations: BA - Brodmann, ASD - autism spectrum disorder, CTL - control or non-autistic individuals, DE - differential gene expression, Ex - Excitatory, IN - interneuron, BBB - Blood brain barrier, ODC - oligodendrocyte, OPC/OPCS - oligodendrocyte precursor cells, UMAP - Uniform manifold of approximation and projection, L23 - cortical layer two and three, L56 - Cortical layer five and six, TF - Transcription factor, Endo – endothelial*

We performed unsupervised clustering of transcriptional signatures from both ASD cases and controls while considering biological and introduced sources of variation (Methods). Visualization of single-nuclei transcriptomes in a uniform manifold approximation and projection (UMAP) grouped nuclei into 35 clusters that were not driven by batch, Brodmann area, or individual samples (Fig. 1B,C; Fig. S1A,B). We annotated the clusters into seven major classes of cortical cell types (neurons: excitatory neurons (Ex neurons), inhibitory neurons (INs) and glial cells: microglia (MG), astrocytes (ASTRO), oligodendrocyte progenitor cells (OPCs), oligodendrocytes (ODC), and blood brain barrier cells (BBB)) based on previously established cell-type-specific canonical gene sets from both human and mouse datasets (15–19), revealing the expected neuronal to glial ratio within human cortex (18) (Fig. 1D; Fig. S1D). Next, we annotated distinct subgroups within each major class by aligning our cell clusters with well-known reference data from the Allen Brain atlas (19), as well as a previous scRNAseq data set from individuals with ASD and controls (13) and cluster-specific differential marker genes (Table S1-3; Methods). All of our clusters paired with the cell profiles previously defined (Fig. 1E; Table S1-2). However, due to the larger number of cells and cohort of individuals profiled here, we define a number of ASD-specific cell states not previously described (Fig. 1E; Table S1-2,4), which mostly represent blood brain barrier cells and reactive glial states that we describe in detail below.

Within non-neuronal populations, BBB subtypes clearly separate based on specific markers into endothelial cells, pericytes, and mural or smooth muscle cell types (20) (Fig. 1D; Fig. S2C). Astrocytes separated into three clusters by expression of specific markers annotating two grey matter/protoplasmic astrocyte clusters, one homeostatic (ASTRO2) and one reactive (ASTRO1), and one white matter astrocyte cluster (21) (ASTRO3; Fig. S2A). We also generated a small spatial transcriptomics dataset (Fig. 1A; Fig. S3; Methods) with sufficient resolution to map specific cellular signatures onto their locations relative to cortical laminae and white matter. To map these cluster specific signatures onto the cortical layers, we filtered cluster marker gene sets into sets of uniquely expressed genes (Methods, Table S1-3). Utilizing this strategy, we confirmed the known patterns of cell composition in the cortex. White matter/fibrous astrocytes are located in white matter and Layer 1, whereas the reactive protoplasmic astrocytes are found within the grey matter layers (22) (Fig. S2H). ODC cluster expression profiles were separated by their maturity and myelination capabilities (ODC1 being the most mature to ODC3, as least mature) (23) (Fig. S2D). OPC clusters are separated based on maturation of myelinating properties and subtype, with OPCS2 expressing the most mature myelination profile and OPCS1 being immature cells, whereas OPCS3 are delineated by *BCAS1* and *GPR17*, which indicate that they are a specific type of OPCS previously found to support synaptic function (24,25) (Fig. S2E). Microglia (MG) subtypes are mainly split by two states: one homeostatic (MG1) and two reactive (MG2 & BBB_MG3) that are differentiated by the expression of known markers (26) (Fig. S2B). We find that homeostatic MG1 are found primarily within white matter and show a sharp drop in abundance within grey matter. MG2 distribution is very similar to homeostatic MG1, but with an increase in abundance in layer 1 (Fig. S2I left and middle panel). BBB_MG3, a rare subclass, are found at uniform but low levels across grey matter laminae in control brains (Fig. S2I right panel).

Ex neuron subtype clusters are separated by expressed known markers associated with their expected superficial-to-deep cortical distribution and axonal projection trajectories (27,28) (Fig. S2F; e.g. *CUX2, LAMB1, RORB, CTIP2/BCL11B*). For Ex neurons, we find that in all cases the cluster expression profiles predict their locations within the spatial transcriptomics data - with layer 2/3 Ex neurons’ marker gene expression located in the superficial layers and Layer 5/6 Ex neurons profiles observed most highly in deeper layers (Fig. S2F). Inhibitory (IN) neuron subtypes are easily annotated based on canonical marker genes and differential gene expression associated with IN ontologies (29–32). We define one somatostatin (SST+) cluster, two parvalbumin+ (PV+) clusters, one vasointestinal peptide cluster (VIP+) that contains calretinin (CR+) cells as well, one reelin (RE+) cluster, and another cluster consisting of KIT+ cells/SV2C+ cells (Fig. S2G).

Mapping the cluster-specific profiles shows that these IN subtypes are distributed evenly across laminae in the spatial data, with the exception of the CGE-derived cluster signature, which are essentially absent in layer 4, matching their known cortical distributions in humans (33).

### Cell Composition changes in ASD

We next perform cell composition analysis to determine if any cell types are reduced or more abundant in the ASD cortex compared to controls (see Methods). In agreement with previous analysis (13,34,35), we detect a slight, but significant, increase in reactive forms of astrocytes and microglia. While previous analyses were limited in sample size and testing methods, our larger dataset affords a higher resolution: we specifically observe an increase in reactive astrocytes (ASTRO1), reactive microglia (MG2) and a trend for an increase in MG3, as well as OPCs (OPCS1 and OPCS3; Fig. 1; Fig. S1G; Table S3-1). Interestingly, when we look at the laminar distribution of the ASTRO1 cluster signature in ASD brains from the spatial transcriptomics data, we find that there is a trend towards increase in ASTRO1 unique marker counts in ASD brains (Fig. 1F; Fig. S2H left panel). This increased trend in reactive astrocyte signature is more pronounced within superficial cortical laminae in ASD relative to deeper layers (Fig. S2H). Likewise, signatures from reactive microglia (MG2) are increased within grey laminae in ASD compared to control (Fig. 1F,Fig. S2I center panel). This tendency for more glial reactivity in superficial laminae and the shift in MG reactivity markers from white matter to grey matter layers is interesting given the previously observed enrichment of ASD risk genes in neurons comprising superficial cortical laminae (3) and the most differential gene expression between controls and ASD occurring in L2/3 Ex neurons (13). We observe no significant changes in neuronal cell type abundance, so we next assessed cell-type transcriptional changes.

### Overview of the cell type specific transcriptional changes in ASD

Not surprisingly, given the large changes observed at the bulk transcriptome level (7,8,14), we detect widespread changes encompassing most cell types, identifying 3485 significant differentially expressed (DE) genes that comprise 2166 down-and 1319 up-regulated genes across 35 cell types (FDR < 0.05;LogFC ≥ 0.2; pseudobulk approach see Methods, Fig. 2B), representing 2810 unique genes, of which only 675 (24%) are DE across more than one cell type. We confirmed subjects with monogenetic form of ASD, (dup)15q11-13, harbor a DE genetic signiature that is highly similar to ASD DE genes within each cell cluster (*R^2^=* 0.86-0.97*, P=*0.0-2.3 × 10 ^-20^ spearman correlation). Notably, out of 2810 DE genes, 333 (12%) genes are known syndromic and Tier one or two ASD risk genes from the SFARI database (OR = 4.1, *P =* 4.9 × 10^-152^, hypergeometric test) and 71 genes are in the syndromic or Tier one SFARI categories, a similar level of enrichment (OR = 1.4 *P =* 7.7 × 10^-29^ hypergeometric test, Methods).

**Figure 2.**
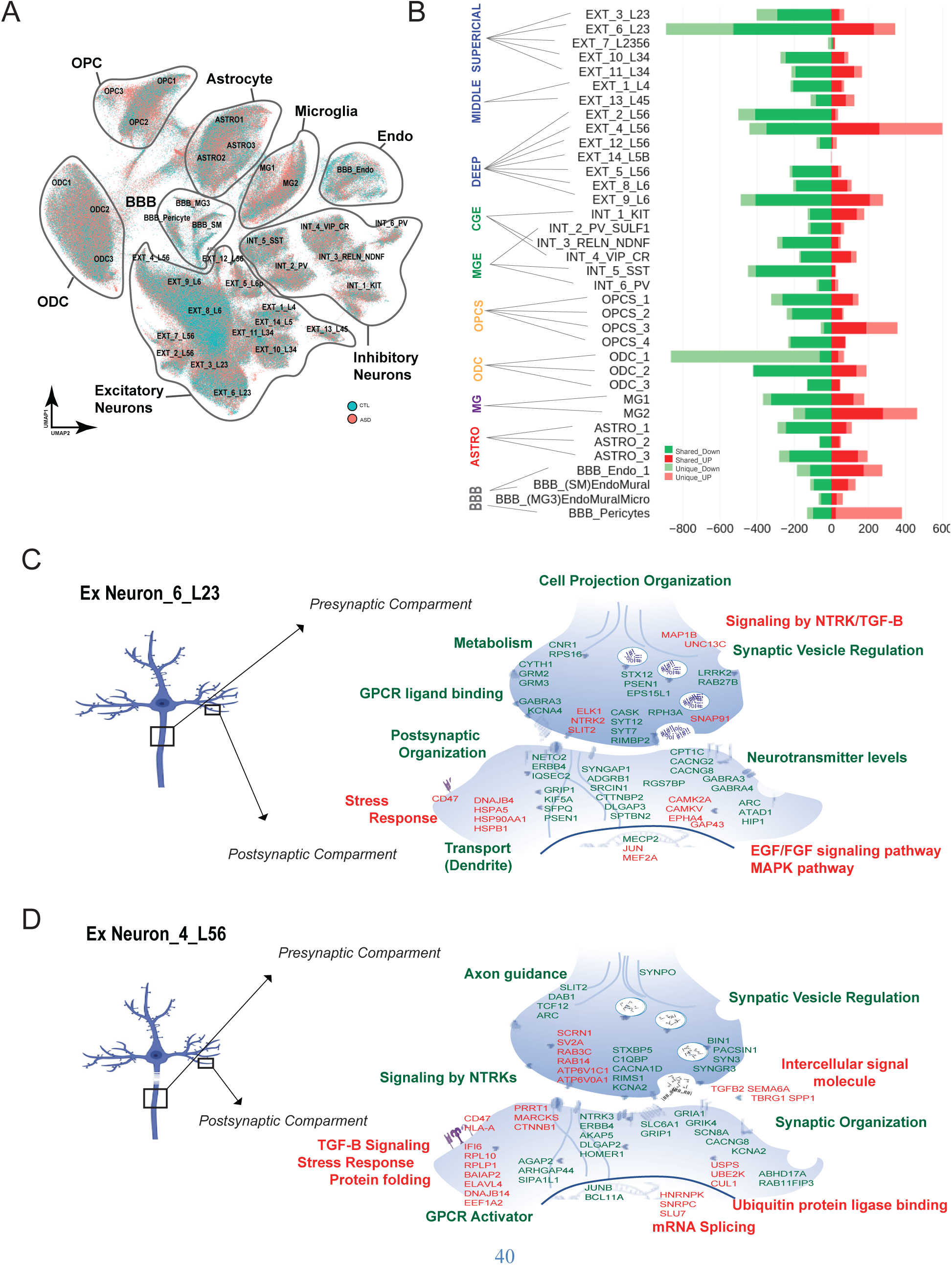
Cell type specific Transcriptional changes. A) UMAP of cell clusters colored by diagnoses status from CTL (blue) and ASD (red) brains, B) Cell type specific count of upregulated genes (red) and down regulated genes (green). Ex neurons are labeled by predicted layer identity (blue), INs are marked by whether they are MGE−(PV+ and SST+ INs) and CGE−(VIP+, CR+, RELN/NDNF+, KIT/SV2C+) derived (green), OPCS and ODCS are labeled in yellow, microglia are labeled in purple and grey, astrocytes are labeled in red and BBB cells are labeled in grey. The number shared genes (solid red or solid green bar) and the number of unique genes (light red or light green bar) vary by cell type. C) Schematic of Ex neuron (blue) cell soma, dendrites, and initial axonal projection, black square indicates putative location enlarged example of the presynaptic axonal compartment on top and enlarged example of postsynaptic dendritic compartment on bottom. The most significant up regulated GO terms and pathways (red) and their accompanying DE gene examples (red); down regulated GO term and pathways (green) and their accompanying DE gene examples (green) are listed on the Ex_6_L23 synaptic components. Membrane-bound receptors and effector molecules may be located on the cell soma as well. D) Synapse schematic of deep layer EXT_4_L56 most significantly up regulated GO terms and pathways (red) and their accompanying DE gene examples (red), down regulated GO term and pathways (green) and their accompanying DE gene examples (green). *Abbreviations: CTL – control individuals, ASD – autism spectrum condition, DE – differential expressed, Ex Neurons – Excitatory Neurons, MGE – Medial ganglion eminence, CGE-caudal ganglionic eminence, PV – parvalbumin or PVALB, SST – somatostatin, VIP – vasointestinal peptide, CR – calretinin, RELN – reeln, GO – Gene ontology, BBB – blood brain barrier, OPCS – oligoprecursor cells, ODC – oligodendrocytes*.

Most changes are cell-type specific, DE genes are also not evenly distributed across cell types. We find that superficial Ex neurons (Ext_6_L23) and reactive glial cell types (ASTRO, MG2 and ODC1) have the highest number of DE genes and the highest number of unique changes in gene expression (e.g. changes not seen in other cell types; Fig. 2B). We also observe high concordance between these transcriptional changes and a previous ASD dataset profiled by bulk RNAseq (7) (R^2^= 0.86*, P=*0; Fig. S1F). Given that approximately half of our samples are not part of any previous bulk tissue and single-cell studies, in part this represents an additional extension and independent replication of that series. Taken together, despite differing approaches, we replicate previously discovered changes, but uncover a greater number of DE genes across cell types. Within neurons, these DE genes are most prominent in specific cell types, highlighting L23 Ex neurons and L56 callosal projecting Ex neurons, which provide long range cortical-cortical connectivity (36,37), and SST INs, which provide local excitatory synaptic balance and modulation of circuit function through inhibitory neurotransmission (38).

### Relationship to previous co-expression modules identified in bulk tissue

We next explore how these single cell expression data were related to known gene co-expression modules differentiating ASD and controls from the two largest and most recent post-mortem, bulk RNAseq studies by identifying significant enrichments of co-expressed module genes within the DE genes of each cell type (Methods; *8, 14*). As expected, a number of modules identified in bulk tissue are non-cell type specific and relate to widespread changes shared across many cell types (Fig. S4A,B; See all modules cell-type associations: Table S4-1), for example modules related to the upregulated stress response and protein translation are enriched across all cells types (modules G-M15 (8), GPC-M27 (14); shared genes include: *EFF1A1, EIF3E, EEF1A1P5, RPL10, RPL27, RPL26*). Pan-neuronal changes occurring across Ex neurons and INs include several downregulated modules related to synaptic, cAMP-, and calcium signaling (GM1, GPC-M19, shared genes: *NRXN2/3, LRRC7, RIMBP2, PLXNA2, DLGAP3, DLG4, CAMK1, SYNGAP1*) (Fig. S4AB, Table S4-1). Pan-glial changes include modules that are enriched for cytokine-mediated signaling, interferon-gamma, and inflammatory JAK/STAT/NFKB signaling that are upregulated most strongly within astrocytes (ASTRO1) and microglia, MG2 and MG3 (Fig. S4A,B; Table S4; G-M5 and GPC-M7, shared genes: *BAG3, BCL6, HLA-E, IFITM2, IRF1/7, NFKBIA, RELA, STAT3, TNFRSF1A*).

We also identify 23 ASD-modules that reveal striking cell-type specificity that were previously inconclusive with respect to their cell type relationships in bulk tissue analysis (8,14) (Fig. S4, Table S4-1). For example, the G-M11 (8) co-expression module is specifically enriched within EXT_6_L23 neurons’ downregulated genes and is enriched in mRNA processing and splicing (Fig. S4A; Table S4-2,3). Likewise, G-M16 is selectively enriched within INT_3_RELN_NDNF interneurons’ downregulated genes, which are enriched in ion channel activity and postsynaptic membrane potential (Fig. S4A; Table S4-2,3). We also implicate G-M26, involving disruption of cell-cell junction assembly and establishment of the BBB (8), which is selectively found in genes downregulated in endothelia (Fig. S4A; Table S4-2,3). The G-M10 module related to unfolded protein response and neuroinflammation was previously not identified as cell type enriched (8). However, the increased resolution afforded by our cell type specific data identifies it as specifically enriched in astrocytes and microglia, as well as ODCS and BBB-related upregulated gene sets (Fig. S4A; Table S4-2,3), which provides new context extending evidence of neuro-inflammation to oligodendrocytes and BBB cells in ASD. Lastly, we find the G-M21 module, which exhibited significant enrichment for Schizophrenia (SCZ) polygenic risk score (39) in previous work (PRS; *8*), to be strongly enriched within the downregulated genes across most Ex neurons, impacting genes involved in transcriptional activity, trans-synaptic signaling, and MAP kinase tyrosine/serine/threonine phosphatase activity (Fig. S4A; Table S4-2,3). These results confirm several previous findings based on network analysis in bulk transcriptome data and meaningfully extend our knowledge of cell-type specificity in brain from individuals with ASD.

### Cell-type specific state changes in ASD

Next, we explore the pathways and mechanisms of cell-type specific transcriptional changes. The most DE genes are detected in superficial Ex neurons (Ext_6_L23), cortical-projecting deep Ex neurons (EX_4_L56), and SST+ interneurons (INT_2_SST), two ASD-specific reactive astrocyte clusters, one protoplasmic and one fibrous (ASTRO1/ASTRO3), reactive MG2, OPCS1 and ODC1(Fig. 2B; Table S5-1).

#### Neuronal changes

Most DE genes in neurons are downregulated and are enriched in GO annotations that converge upon synaptic components, both postsynaptic and presynaptic, as exemplified by changes in L23 Ex neurons, L56 Ex neurons, and SST+ IN (Fig. 2C,D; Table S5-1; see also Fig. S5A,B; Methods). For superficial ex neurons (EXT_6_L23), the most significant pathways downregulated are related to dendrite/cell projection (axon) organization (e.g. *NETO2, GRIP1, DLGAP3, ARC, ATAD1*), vesicle regulation (e.g. *HIP1, LRRK2, RAB27B, STX12, CYTH1*), neurotransmission (e.g. *GABRA4, CACNAG2, CACNG8, RIMBP2, SYT12, CNR1*) and GPCR ligand binding (e.g. *GRM2, GRM3, IQSEC2, ERBB4*; Fig. 2C; Table S5-1; Table S6A). The upregulated pathways in these cells include oxidative phosphorylation and inflammatory stress responses (e.g. *RPS16, CD47, DNAJB4, HSPA5, HSP90AA1, HSPB1, JUN),* signaling by NTRK and TGFB receptors (e.g. *NTRK2, SLIT2, MAP1B, UNC13C, JUN, FOS*), and growth factor response TGFB/EGF/MAPK pathways (e.g. *CAMK2A, CAMKV, EPHA4, GAP43*) (Fig. 2C; Table S5-1). In contrast to EXT_6 L23 neurons, EXT_4_L56 cortical-projecting neurons manifest downregulation of signaling by NTRKs (e.g. *NTRK3, AKAP5, DLGAP2, SLIT2*) and neurodevelopmental genes related to axon guidance (e.g. *DAB1, ARC, SYNPO, TCF12*; Fig. 2D, Table S5-1). These neurons also exhibit upregulation of genes related to mRNA splicing (e.g. *HNRNPK, SNRPC, SLU7*), ubiquitin protein ligand (e.g. *USPS, UBE2K, CUL1*), a stronger inflammatory response (e.g. *CD47, HLA-A, IFI6, RPL10, RPLP1, EEF1A2, DNAJB14*), and intracellular signaling involving pro-inflammatory/TGFB signals (e.g. *TGFB2, SEMA6A,* and *SPP1*; Fig. 2D; Table S5-1; Table S6A). These data provide evidence that two classes of excitatory intercortical projection neurons exhibit the most profound transcriptional dysregulation in ASD, implicating disruption in long-range cortical connectivity. Both populations significantly up-regulate pro-inflammatory signals. Yet, these two clusters of Ex projection neurons in different cortical laminae display distinct patterns of down-regulation in genes comprising biological pathways that are broadly related to neuronal signaling and neurodevelopment.

We next compare the transcriptional changes in both Ex cell types highlighted above to changes within INs showing the largest DE changes, the SST+ INs (INT_5_SST). Here, we observed a similar downregulation of pathways related to synaptic vesicle regulation, neurotransmission, and synaptic organization (Fig. S5A; Table S6A). However, INT_5_SST neurons showed a more striking downregulation of genes directly related to glutamatergic receptor function and signaling (e.g. *GRIA1/2/3, GRIN2A/3A, GRIN2B, HOMER1, SHANNK3, NLGN1, TIAM1*), structural and functional components of the action potential (e.g. *SCN3B, ANK2, SCN1A, SCN8A, SCN2A)* and mRNA splicing, including members of the RBFOX and STAR families (e.g. *RBFOX1, SLM1, SLM2*), which have been shown to play critical roles in synaptic connectivity in these cells in mouse (40,41) (Fig. S5A). The upregulated genes in SST+ INs overlapped with several pathways up-regulated in EXT_4_L56 cortical projection neurons including increases in inflammatory response (e.g. *HLA-A, B2M, IGF1R)* and protein modification changes, as well as an increase in transport along cytoskeleton and calcium mediated axon guidance (e.g. *RAC1, RDX, SCNA, MAPT*; Fig. S5A; Table S6A). Mapping the SST+ IN cluster-specific unique downregulated DE genes onto the spatial transcriptomic dataset demonstrates that this DE signature was most prominent within superficial layers in the CTL cortex, suggesting that SST+ Ins in these laminae are relatively the most affected (Fig. S5A; right panel, *P = 0.007* in L2/3, linear mixed model, Methods).

#### Glial changes

Glial cells in ASD are primarily characterized by changes to the state of these cells from a homeostatic, or resting state, to reactive or activated inflammatory states, compared with controls. This change is illustrated by upregulation of cell-type specific responses to inflammatory and interferon signaling and a parallel decrease in expression of homeostatic genes (21,26). We describe below the ASD reactive inflammatory states based on the major glial types: Astrocytes, Microglia and ODCs.

Genes upregulated in protoplasmic (PP), grey matter astrocytes (ASTRO1) represent astrocyte proliferation (e.g. *FOXO3, RFX7, ZEB1*) that likely lies downstream from TGF-B/ interferon signaling through MHC and other neural-immune receptors (e.g. *HLA-B, HLA-C, HLA-E, CD44, IFI6, IFITM2, IFGR1, TGFBR2, BMP7*) and their inflammatory response effectors (e.g. *IRF1, JAK3, MED12L, SMARCA2, STAT1/3, SMAD1/3*; Fig. 3A; Table S6B) (42,43), which are upregulated in parallel. Fibrous (FB) astrocytes (ASTRO3) exhibit similar changes, such as upregulation of MHC and other receptors (e.g. *HLA-B, HLA-C, HLA-E, CD99, CD44, NTRK2*), but differ by upregulation of cytoskeleton morphogenesis and focal adhesion signaling (e.g. *ITGB5, ITGAV, TAB2, KALRN, NAV3*), suggesting changes in morphology or projections from white matter FB-astrocytes in cortex in individuals with ASD. FB-astrocytes (ASTRO3) also upregulate genes related to stress-related apoptosis and unfolded protein response (e.g. *DNAJA4, HSP90AA1, HSPA1B, BAG3, DAB2IP*) (Fig. 3B; Table S5-1). Both PP-and FB-astrocytes exhibit a downregulation of lipid and cholesterol biosynthesis pathway genes (e.g. *HMGCS1*). However, PP-astrocytes possess downregulated growth factor and NTRK signaling genes, whereas the opposite is true for FB-astrocytes in the ASD cortex samples relative to controls. Specifically, FB-astrocytes upregulate expression of TGFB signaling (e.g. *TGFB1*). growth factors (e.g. *VEGFA*) and extracellular matrix proteins (e.g. *ADAM12, ADAMTS17*) (Fig. 3B; Table S6B).

**Figure 3.**
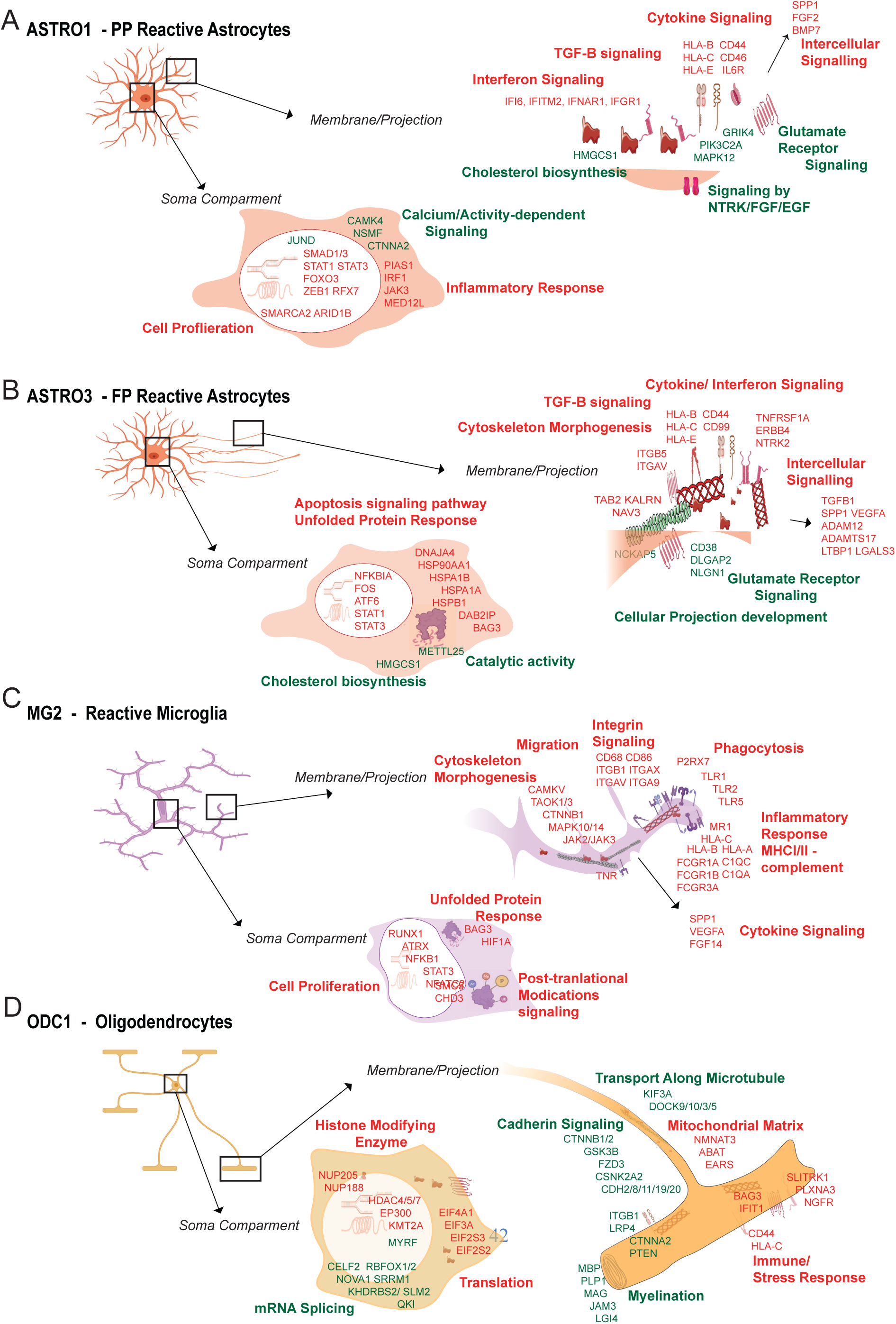
Glial cell-type specific transcriptional changes. A) Cartoon of PP reactive astrocyte (orange), ASTRO1; black box indicates putative location of the DE genes in enlarged astrocyte membrane/ projection, top, and soma compartment, bottom. We illustrate the most significant up regulated GO terms and pathways (red) and their accompanying DE genes (red), down regulated GO term and pathways (green) and their accompanying DE gene examples (green). B) Cartoon schematic of WM reactive astrocyte (orange), ASTRO3. The most significant up regulated GO terms and pathways (red) and their accompanying DE gene examples (red), down regulated GO term and pathways (green) and their accompanying DE gene examples (green). C) Cartoon of reactive MG2, the most significant up regulated GO terms and pathways (red) and their accompanying DE gene examples (red). D) Schematic of oligodendrocytes, ODC1, with a black box indicating putative location of the DE genes in long enlarged myelinating membrane/ projection, top, and soma compartment, bottom. The most significant up regulated GO terms and pathways (red) and their accompanying DE gene examples (red), down regulated GO term and pathways (green) and their accompanying DE gene examples (green) are presented. *Abbreviations: PP-protoplasmic, DE-differentially expressed, GO – Gene ontology, WM – White matter, MG – microglia, ODC - olidodendrocyte*

Microglia (MG2) manifest an increase in reactive state markers highlighted by the up-regulation of known pro-phagocytotic genes (e.g. *P2RX7, TLR1/2/5, CD68*) including pronounced up-regulation of the extracellular signaling molecule SPP1 (44) (Fig. 3C). MG2 cells also express a higher number of MHC I/II class molecules, complement, and receptors (e.g. *HLA-A/B/C, FCGR1A/1B/3A, C1QC, C1QA, TNR, TGFBR1*), more so than the other glial cell classes. In ASD, reactive MG2 cells also increase genes related to cell morphogenesis (e.g. *CAMKV, TAOK1/3, CTNNB1, MAPK10/14, JAK2/3*) and integrin signaling (e.g. *CD68, CD86, ITGB1, ITGAX, ITGAV, ITGA9*), which suggests an increase in migration and formation of contacts with other cells (Fig. 3C, Table S5-1; Table S6B). Since many of these changes are downstream of cytokine signaling, we hypothesize from these data that SPP1 may be acting as an extracellular element responsible for coordinating the cell-cell interactions putatively directing synaptic and/or cellular removal in activated MG (44,45).

Interestingly, we observe that ODCs are also shifted from a homeostatic state to a more immuno-reactive state, which has not been previously observed in ASD. For example, the oligodendrocyte cluster, ODC1, upregulates genes related to inflammatory and energy stress response (e.g. *IFIT1, BAG3, ABAT, EARS2, NMNAT3, EIF4A1, EIF3A*) and histone modifying enzymes (e.g. *HDAC4/5/7, KDM5A, EP300, KMT2A*) (Fig. 3D, Table S5-1). However, ODC1 cells also have a substantial number of downregulated genes, the largest of any glia cell type and similar in magnitude to that of superficial L23 Ex neurons (Fig. 2B). Enriched down-regulated pathways include cadherin signaling (e.g. *CTNNB1/2, GSK3B, FZD3, CDH2/8/11/19/20*), transport along microtubule (e.g. *KIF3A, DOCK9/10/3/5*) and myelination (e.g. *MBP, PLP1, MAG, JAM3, LGI4, MYRF*), which predict a potential reduction in ODC1’s capabilities to mature and functionally extend their process, adhere and myelinate neuronal axons properly (Fig. 3D; Table S5-1; Table S6B).

### Defining the regulatory networks underlying transcriptional changes

We next systematically integrate the cell-type specific ASD DE data with transcription factor (TF) networks to discover candidate drivers of the observed transcriptional changes in ASD relative to controls and to determine if these could be related to causal genetic factors (Fig. 4A). We analyze the single cell transcriptomes with SCENIC (46,47), which robustly identifies TF networks driving cell-type specific changes (Fig. 4A). The networks, called regulons, consist of the TF and its down-stream target genes (that are significantly co-expressed and harbor the TFs motif within 10kb of TSS), which we then score to identify which are preferentially associated with ASD followed by their overlap with cell-type specific DE genes (Fig. 4A; Methods). We identify 217 of 337 total regulons detected that are predicted to be changed in ASD in at least one cell type (Table S8-1,2). We summarize the top significant regulons predicted to regulate the most DE genes by cell type, L23 Ex neurons, SST+ INs, and MG2 (Fig. 4B,C, Methods), finding that these regulons can account for an impressive proportion of the observed differential gene expression (Fig. 4D-F; Table S8-5,6,7). For example, we found *CUX1* regulons to be a major driver of DE across cell types: CUX1 is predicted to regulate 40% of downregulated genes within L23 Ex neurons and 44% of downregulated genes with SST+ INs (Fig. 4B,D,E). Additionally, if we consider the top 6 regulons overlap with downregulated genes in each neuronal cell type, we find that collectively they are predicted to regulate 68% of the downregulated genes within L23 Ex neurons (e.g. *CUX1, CHD2, ZNF91, BDP1, SREBF2, KDM5A* regulons) and 75% of downregulated genes within SST+ INs (e.g. *CUX1, CHD2, ZNF91, SREBF2, ZEB1, SOX6* regulons; Fig. 4D,E). The CUX1 downregulated networks within both neuronal types form significant direct protein-protein interaction (PPI) network, and are significantly enriched for ASD-associated risk genes including a number of TFs that are also regulon drivers (e.g. TAF1, SREBF2, KDM5A, BCL11A; Fig. 5A,B; Methods). The *CUX1* regulon within EXT_6_L23 are genes that function within the synapse, Golgi apparatus and nucleus, related to catalytic activity on a protein (e.g. *SYNJ2, MTOR, CYTH1, PSEN1, PLCB4, IQSEC2, ERBB4, SFPQ, HIP1*) and gene expression (e.g. *EP300, MED27, SMAD3, SREBF2, NPAS2, NCOA3, NCOR2*; Fig 5A). The *CUX1* regulon set of downregulated genes within SST+ INs is enriched for genes that function in synaptic signaling, such as calcium and potassium voltage-gated ion channels (e.g. *CACNA1B, CACNA1A, KCNQ3, CACNA1D, KCNC2, CACNA1C*) and GTPase activity (e.g. *TIAM1, RASGRF2, FAM49B, KALRN, AUTS2, STMN*1; Fig. 5B,D).

**Figure 4.**
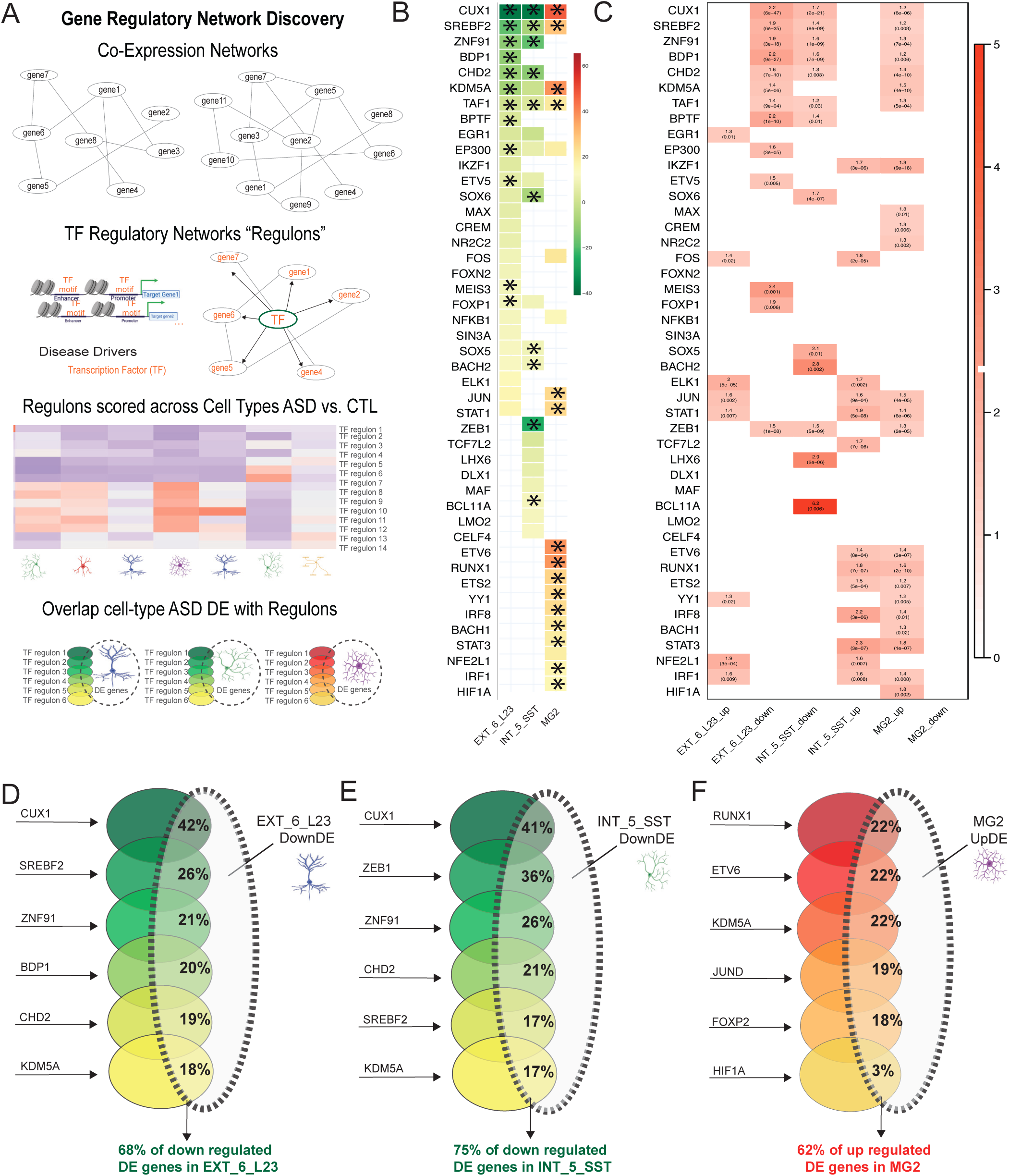
Regulatory networks driving cell-type specific changes in DE genes. A) Schematic of network discovery using SCENIC; initially co-expression networks are generated from cellular transcriptional profiles then pruned to TF-driven regulatory networks, called ‘regulons’, consisting of the TF and its down-stream target genes (that are significantly co-expressed and harbor the TFs motif within 10kb of TSS); regulons are then scored to identify which are changed within ASD followed by assessing their overlap with cell-type specific DE genes B) Overlap percentage of TF-regulon genes with EXT_6_L23, INT_5_SST INs, and MG2 ASD DE genes with downregulated DE genes (green) and upregulated genes (red). Black stars indicate significant overlaps in shown in (C), C) Heatmap of odd ratios (OR, red) between regulons separated by upregulated and downregulated DE genes within EXT_6_L23, INT_5_SST INs, and MG2 cell types. Associated OR and p-values are printed in black over each significant association. D) Overlap percentage between EXT_6_L23 down regulated DE with top regulons predicted to regulate DE genes (green to yellow). These regulons together account for 68% of downregulated DE in EXT_6_L23. E) Overlap of INT_5_SST down regulated DE with top regulons predicted to regulate DE genes (green to yellow). Together they account for 75% of downregulated DE genes in SST+ INs. F) Percent overlap MG2 up regulated DE with top regulons (red to yellow). Together they account for 62% of upregulated genes in MG2 cells. *Abbreviations: TF – transcription factor, ASD – autism spectrum condition, DE – differentially expressed, OR – odds ratio, EXT – excitatory neuron cluster, INT/INs – interneuron cluster*.

**Figure 5.**
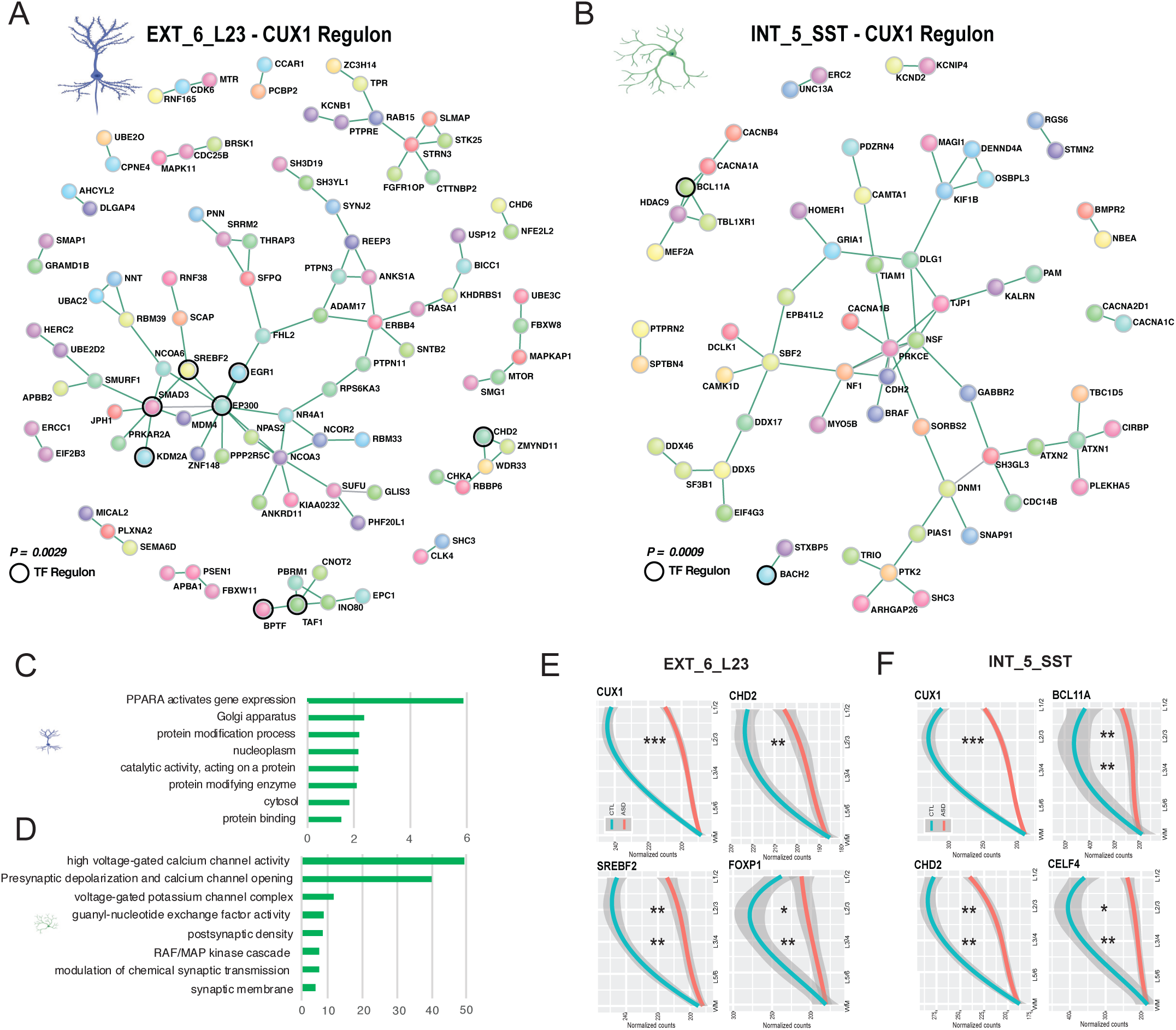
*CUX1* gene network in neurons forms significant PPI network containing ASD risk genes and regulon TF drivers. A) PPI network formed from the *CUX1* down regulated DE genes in EXT_6_L23 cells (*P = 0.0029,* 1000 permutations) illustrating a striking abundance of ASD risk genes and other TF-drivers of regulons (black outlined circles) predicted to interact. B) PPI network formed from the *CUX1* downregulated DE genes specifically in INT_5_SST cells (*P = 0.0009,* 1000 permutations) illustrating a similar abundance genes of ASD risk genes and other TF-drivers of regulons (black outlined circles) predicted to interact. C) Significant enrichment of GO terms (green) of the downregulated *CUX1* regulon genes within EXT_6_L23 DE. Genes from all GO categories, E) Significant enrichment of GO terms (green) of the downregulated *CUX1* regulon genes within INT_5_SST from all GO categories, E) normalized count of EXT_6_L23 down regulated genes that overlap top regulons *CUX1, CHD2, SREBF2,* and *FOXP1* are located highly in superficial layers of CTL cortex (blue line) and reduced most significantly in the ASD superficial layers as expected (red line), F) normalized count of INT_5_SST down regulated DE genes that overlap top regulons *CUX1, CHD2, BCL11A*, and *CELF4* regulons are found located highly in superficial layers of CTL cortex (blue line) and reduced most significantly in the ASD superficial layers (red line). *0.01<*P*<0.05 **0.001<*P*<0.01 \*\*\**P*<0.001. *Abbreviations: PPI – protein-protein interaction network, DE – differentially expressed, TF-transcription factors, P – Pvalue, CTL-control individuals, ASD – autism spectrum condition*.

The spatial distributions of down-regulated genes from the L23 Ex neuron and SST+ IN regulons indicate that these networks are most reduced in the ASD superficial layers 2/3 (Fig. 5E,F; EXT_6_L23 regulons: *CUX1* regulon down DE genes Layer × Condition interaction, *P= 1.01×10^-142^*; L2/3 ASD vs. CTL, *P= 8.92 x10^-05^*; *CHD2* Layer × Condition interaction, *P= 1.40×10^-30^*, Layer 2/3 ASD vs. CTL *P= 0.002*; *SREBF2* Layer × Condition interaction, *P= 1.55×10^-44^*; Layer 3/4 ASD vs. CTL *P= 0.0003*; Layer 2/3 ASD vs. CTL, *P= 0.0004*; *FOXP1* Layer × Condition interaction *P= 6.61×10^-12^*, Layer 3/4 ASD vs CTL, *P = 0.004*; Layer 2/3 ASD vs CTL *P = 0.013*; INT_5_SST regulons: *CUX1* Layer × Condition interaction *P= 9.11×10^-08^*; L3/4 ASD vs CTL *P= 0.0002*; Layer 2/3 ASD vs. CTL *P = 0.0002*; CHD2 Layer × Condition interaction *P= 6.63 × 10^-66^*; Layer 3/4 ASD vs. CTL, *P= 0.001*, Layer 2/3 ASD vs. CTL *P = 0.001*; BCL11A Layer × Condition interaction *P= 4.76 x10^-12^*, Layer 3/4 ASD vs. CTL *P= 0.001*, Layer 2/3 ASD vs. CTL *P = 0.006*; CELF4 Layer × Condition interaction *P= 9.16 x10^-11^*; Layer 3/4 ASD vs CTL *P = 0.008*; Layer 2/3 ASD vs. CTL *P = 0.021*; FOXP1 Layer × Condition interaction *P= 4.69 × 10^-07^*; Layer 3/4 ASD vs. CTL *P = 0.0062*; Layer 2/3 ASD vs. CTL *P = 0.0093*, linear mixed model, Methods).

We summarize the top regulons identified in reactive MG2 in ASD, which also account for a significant proportion of DE genes (Fig.4B,C,F; Table S8-1,6); the top 6 regulons account for up to 62% of up regulated DE genes (Fig. 4F). Of note, we find that *RUNX1* and *ETV6* regulons are activated in MG2 and account for 21%, and 22%, respectively, of upregulated genes within MG2 in ASD (Fig. 4B,C,F). The *RUNX1* regulon forms a significant, small PPI network including a few other regulon drivers (e.g. *KDM5A, RUNX2*; Fig. S6A). The MG2 *RUNX1* regulon includes up-regulated genes that act in the complement cascade, phagocytosis, and stress response (e.g. *TLR1/2/5, FCGR1A, HIF1A, BCL6*) and also a significant proportion of down-regulated genes related to MG homeostasis (e.g. *CX3CR1, CSF1R, LTC4S, PDE3B, SLCO2B1, ADGRG1*; Fig. S6A,B; Table S8-1,6). Interestingly, *RUNX1* is induced by TGFB signaling (48) - *TGFB1* is increased in FB reactive astrocytes (Fig. 3B) and *TGFB2* is increased in L56 Ex neurons (Fig. 2B). TGFB receptors are also increased in MG (Fig. 3C), most other neurons (Table S6A) and *SPP1* expression is increased across these same cell types, which is predicted to promote pro-inflammatory, pro-phagocytic neuron-microglia interactions (45). Moreover, TGFB1 signaling has been show to activate *JUN/FOS* in neurons as part of the cytokine inflammatory pathway (49)– consistent with this, we observe *JUN/FOS* expression and other pathways associated with TGFB signaling increased in neurons as well (Fig. 2C,D; Fig. 3; Fig. S5; Table S6A, Table S6B).

#### TF foot-printing to validate regulatory network predictions

Although SCENIC predictions have been shown to be highly reliable in predicting experimentally validated regulatory networks (1,50–52), they are computational. So, we experimentally validate key regulatory changes by performing snATACseq from a subset of the same 22 samples (12 cases and 10 controls) subjected to scRNAseq (Methods; Table S1-6). We generated over 100,000 sn profiles after quality control filtering, which we integrate with the snRNAseq to verify and validate cell-type cluster annotations (Fig. 6A; Fig. S7A; Methods). We find a high concordance between the snATAC and snRNAseq clusters delineated by high correlations between cluster profiles (Fig. S7B; Methods).

**Figure 6.**
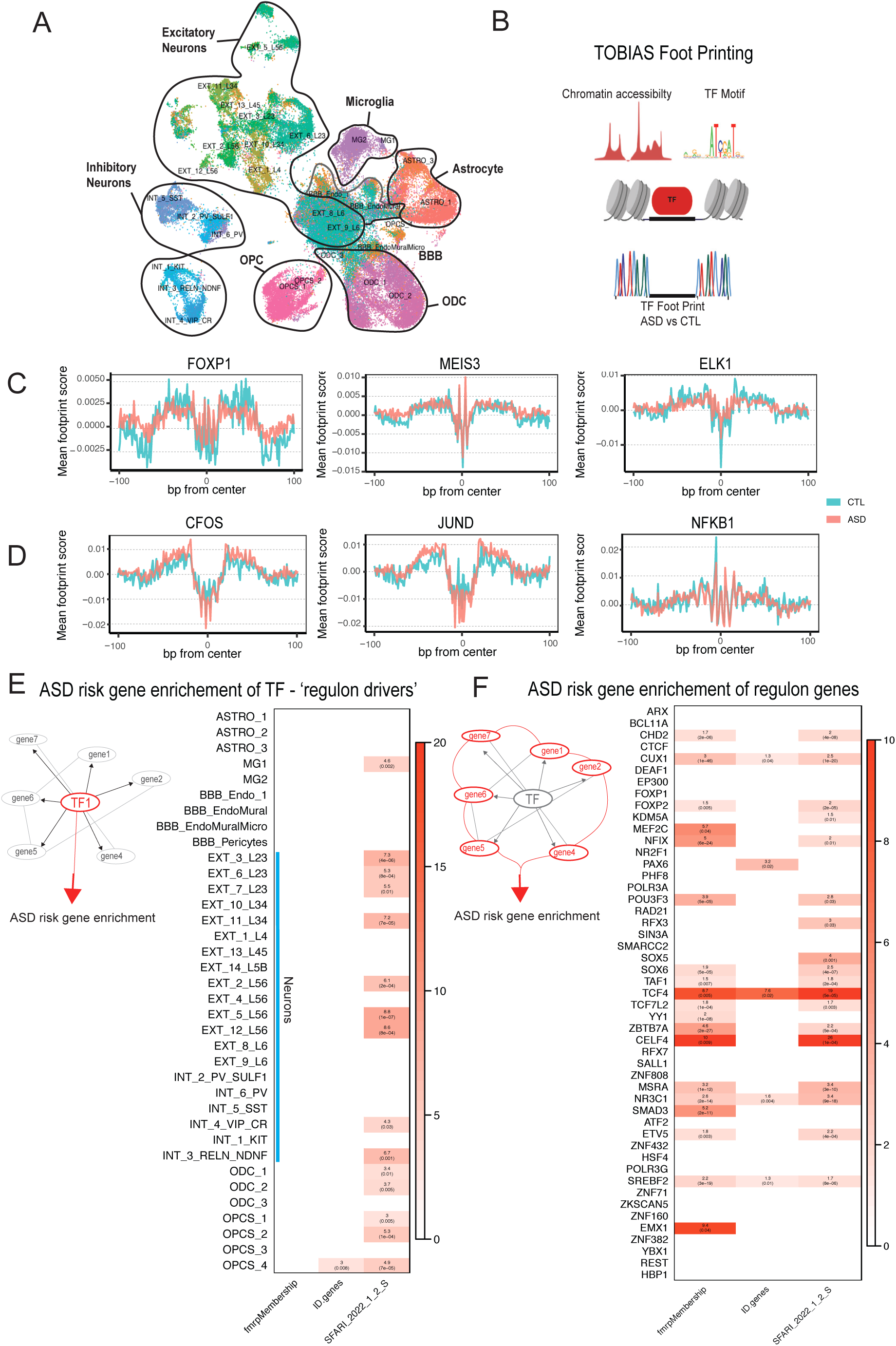
Neuronal cell classes and homeostatic glia types are enriched for regulons driven by ASD risk TF genes. A) UMAP of cellular clusters from scATACseq computationally annotated by scRNAseq clusters, B) Foot-printing approach with TOBIAS to assess changes in TF occupancy within open chromatin profiles of each cell type. C) Representative TF foot-print of significantly downregulated TF occupancy in open chromatin of aggregated Ex Neurons from CTL (blue trace) and ASD (red trace) brains for *FOXP1* (left panel), *MEIS3* (middle panel) and *ELK1* (right panel). D) Representative TF foot-print of significantly upregulated TF occupancy in open chromatin of aggregated MG cells from CTL (blue trace) and ASD (red trace) brains for *CFOS* (left panel), *JUND* (middle panel) and *NFKB1* (right panel). E) ASD risk enrichment heatmap of TF-drivers by cell type, OR (red) and *P*-value included (black) for each cell type, neuronal clusters marked by blue line. F) Risk gene enrichment heatmap of regulon genes to gene sets associated with FMRP, ID, and ASD risk genes, OR (red) and *P*-value for each regulon. *Abbreviations: TF – transcription factor, Ex Neurons – Excitatory Neurons, MG – microglia, CTL – control individuals, ASD – autism spectrum condition, FMRP – Fragile × syndrome polyribosome-associated neuronal RNA-binding protein, ID – Intellectual disabled, OR – odds ratio*.

We next utilize the open chromatin profiles from each major cell type to assess changes in transcription factor foot-printing using TOBIAS (53,54) (Fig. 6B; Methods), focusing on TFs predicted to account for the major changes in DE described above (Fig. 2–3; Fig. S5). Indeed, within Ex neurons and INs we observe that many of the TFs predicted to drive downstream DE genes within the network have changes in occupancy within the relevant DNA foot-prints (e.g. *CUX1, CHD2, SREBF2*; Fig. S8D,E; Fig6C,D). For example, in L23 Ex neurons we detect significantly reduced TF activity scores within a number of networks such as *FOXP1*, *MEIS3* and *ELK1* (Fig. 6C, Table S10; Fig. S7D). We also see increased TF activity for activated pathways in neurons, for example *EGR1* and *JUNB* (Fig. S7D,E; Table S10; Fig. 6D). Likewise within reactive MG2, we see increases in TF activity for *IRF1, IRF8, JUND/B, MAF*, and *EP300* (Fig. S7F, Table S11) with examples of significantly altered foot-prints for *CFOS*, *JUND*, and *NFKB1* (Fig. 6D). Thus, together, we find the many of the predicted changes from the SCENIC regulatory networks and DE data are supported by TF occupancy changes within open chromatin in ASD relative to controls.

### ASD risk gene enrichment

To determine whether these regulons represent known causal aspects of ASD biology, we next test whether the drivers or their downstream DE targets are enriched in known ASD risk genes (Syndromic, Tier one and two; Methods). Indeed, 30 of the regulons dysregulated in ASD brain are driven by high-confidence TF risk genes from the SFARI database, a significant enrichment (e.g. *CUX1, CHD2, FOXP1, BCL11A, SOX5*, Fig. S8A, *OR=1.64, P= 0.004*, hypergeometric test). MEF2C, one of the TF drivers of regulons in Ex neurons, was identified previously as a point of convergence of ASD risk genes based on co-expression analysis in developing brain (3). Further, 21 of these regulons are enriched for association with common ASD PRS (55), a highly significant over-representation (e.g. *CELF4, SREBF2, ETV5, NR3C1*; *P = 0.049-0.0007*, Methods; Table S8-4; Fig. S8A). These regulons are primarily DE within Ex and IN neuron populations (highest in Ex neurons) and less so in glia, consistent with previous studies of ASD risk gene expression patterns (50,56) (Fig. 6E,Fig. S8A). However, we note that there are several TF risk regulons (e.g. *SOX5, SOX6, TCF7L2, PAX6, SREBF2*) differentially expressed in ASD ASTRO1, homeostatic MG1, and OPCS, as well as neurons (Fig. S8A). ASD risk enrichment has not been demonstrated before in glial cells. These regulons comprise genes that are down-regulated, which mechanistically is the expected direction, since the observed down-regulation of TF drivers is the expected scenario occurring in haploinsufficiency of a risk gene, which is predicted for most of these genes in ASD. We also examine whether these regulons’ target genes overall are enriched for ASD risk genes (SFARI tier 1, 2 or syndromic) and genes associated with intellectual disability (ID) (57) and FRMP targets (58). Indeed, many regulons driven by ASD TF risk genes or those enriched for ASD associated common genetic variation are also significantly enriched for ASD risk genes ( n = 19) and FRMP targets (n = 17), while only a few are enriched for ID risk genes (n = 5; Fig. 6F, Fig. S9) in comparison to other regulons. Taken together, these represent a critical starting point to understanding the mechanistic changes driven by risk genes in the brains of individuals with ASD.

## Discussion

We perform the most comprehensive single cell genomic assessment of ASD to date, extending previous observations and providing new insights into ASD biology. The markedly increased depth and breadth of this single cell analysis, involving double the number of individuals and eight times the number of cells profiled, solidifies our picture of the major cortical cell types impacted in ASD, including alterations in cell type composition, remodeling of cell states and the identification of cell-type specific transcriptional cascades. The integration of snRNAseq, snATAC-seq and layer specific spatial transcriptomics allows us to describe and validate candidate regulatory networks driving cell type specific transcriptional changes and their location within cortical laminae. Although we observe broad transcriptional changes in both inhibitory and excitatory neurons, consistent with previous observations in bulk tissue, the magnitude of changes varies considerably by cell type. We find that the most changes in neuronal gene expression occur in glutamatergic neurons within the superficial layers of the cortex, confirming a previous analysis implicating superficial layer projection neurons (13), and further identifying large changes in somatostatin interneurons and cortical-cortical projecting L56 neurons. Moreover, these changes in neuronal gene expression in ASD are accompanied by marked shifts to more activated pro-inflammatory states in multiple glial cell types, including ODCs, MG, astrocytes and cells that comprise the BBB.

It is well recognized that transcriptomic changes associated with a neuropsychiatric condition, although representing a quantitative genome wide molecular measurement, may represent the consequence of that condition, and do not necessarily imply causality. We reasoned that by identifying potential transcriptional regulators and subsequently, to test whether these TFs or their downstream targets were enriched in genetic risk, that we could implicate potential causal drivers. Our analysis identifies candidate TF drivers that account for a substantial portion of major gene expression changes in multiple neuronal and glial cell types, many of which were validated by TF foot-printing. That these TFs coupled with their down downstream targets, termed regulons, are enriched in known high confidence ASD risk genes harboring rare variants and common polygenic risk for ASD, supports their potential causal role in ASD pathogenesis. Moreover, these analyses delineate the potential down-stream consequences of these causal genetic variants in ASD brain by identifying differentially expressed genes and the biological pathways that are regulated by these risk genes in human brain. These regulon networks associated with ASD risk reflect downregulation of synaptic vesicle regulation, neurotransmission, and components related to synaptic development and organization pathways, which have been observed in previous work in other samples (8,13,14). Interestingly a number of these TFs have important roles in the development and maintenance of cell identity (ex, *CUX1, SOX6, FOXP1*), or are markers for neuronal types or cortical laminae, providing a link between fetal development (e.g. ASD loci enrichment in specific TFs) and its postnatal ramifications (cortical transcriptional changes in ASD). We emphasize that more definitive determination of the causal impact of risk genes on these transcriptional regulatory circuits awaits direct experimental perturbations, for example in human *in vitro* models (59–61).

These data support previous predictions of ASD risk gene enrichment in superficial layer neurons from developing neurotypical human cortex (3,62) and significantly extends these observations by: a) implicating superficial SST+ INs and deep-layer, interhemispherically-projecting Ex neurons; b) linking regulatory networks driven by ASD risk-TFs to cell-type specific DE genes in postnatal brain in ASD; c) showing that the shift in expression of homeostatic genes to genes that underlie highly ‘reactive’ or ‘activated inflammatory’ state in response to their environment in microglia and astrocytes, also extends to ODCs and cells that comprise the BBB; 4) and that some of these changes in microglia and ODCs, are linked to multiple candidate TF drivers that are known ASD risk genes, providing a potentially causal link to some of the reactive changes in these glial cells. Further, these data highlight the potential downstream consequences of many of the TF drivers, which act primarily in neurons, and that may subsequently influence glial reactive states, since the latter are also more concentrated in superficial cortical laminae, where both risk genes and transcriptional changes are most prominent. Recent work has demonstrated cortex-wide transcriptional changes in ASD brain (14), including primary sensory regions, which is consistent with the preferential involvement of neurons responsible for cortical-cortical and interhemispheric connectivity. These data provide a starting point for developing mechanistic, molecular and cellular explanations for observations of changes in short and long-range cortical-cortical connectivity and cortical circuits in ASD (3,63–65).

Autistic individuals have a diverse array of co-existing conditions (i.e. epilepsy, attention deficient hyperactivity disorder, schizophrenia, gastro-intestinal conditions, cerebral palsy, bipolar disorder), traits (i.e. apraxia or speech difficulties, social communication differences, sensory sensitivities, learning differences, specialized interests), and related behaviors (i.e. repetitive movements or “stimulatory-seeking”, echolalia, etc.) that make up their individual characteristics, or ‘neurotype’ of ASD (55,66–70). Thus, it will require large numbers of individual samples to elucidate sub-groups and to definitively match specific patient traits, such as cognition or social behavior, to the observed molecular and cellular changes observed here. We note that we included tissue from five individuals with a monogenic form of ASD, (dup)15q11-13 in this analysis, which have previously been demonstrated to share the major transcriptional alterations and pathways observed in idiopathic ASD at the bulk level, which we observe here at the single cell level, as well. A major challenge is now to understand how these cell-cell interactions develop and evolve at both a sub-cellular and circuit level and to determine how they relate to physiological and behavioral states.

## Methods

### Sample Acquisition and preparation for single-nuclei RNAseq and snATACseq

Postmortem cortical brain samples were acquired from the Autism Brain Net project (ABN, formerly the Autism Tissue Project at Harvard, ATP) and the University of Maryland Brain Banks (UMDB) through the NIH NeuroBioBank. A total of 64 samples from subjects with ASD, dup15q syndrome, and non-psychiatric controls from the prefrontal cortex (PFC) or from Broadman area 9 (BA9) and to a smaller extent BA4/6. Thirty one subjects had not been previously profiled. BA4/6 samples were previously subjected to bulk studies: Parikshak et al., *Nature* 2016 (7); Gandal et al., *Science* 2018 (8) and a small subset of samples (6 samples) were processed for a different single-cell analysis in Gandal et al., *Nature* 2022 (14).

Frozen brain samples were placed on dry ice in a dehydrated dissection chamber to reduce degradation effects from sample thawing and/or humidity. Approximately 20-50mg of cortex was sectioned, ensuring specific grey matter to white matter boundary and ratio. The tissue section was homogenized in RNase free conditions with a light detergent (0.001% Triton100X/PBS/1%BSA/RNase) briefly on ice using a dounce homogenizer, filtered through a 40uM filter and centrifuged at 1000 g for 8 minutes at 4C. The pelleted nuclei were then filtered through a two-part micro gradient (30%/50% Iodixanol Sucrose medium) for 20 mins at 4C. Clean nuclei were pelleted away from debris. The nuclei were washed two more times with PBS/1%BSA/RNase and spun down at 500g for 5 mins. Cells were inspected for quality (shape, color, membrane integrity) and counted on a countess II machine. They were then loaded onto the 10X Genomics platform to isolate single nuclei and generate libraries for RNA sequencing on the NovaS4 or NovaS2 Illumina machines for a depth of about 65K reads per cell.

### snRNAseq Data Processing

After sequencing, Cell Ranger software from 10X Genomics (https://support.10xgenomics.com/single-cell-gene-expression/software) was used to prepare fastq file and reads were aligned to the human GRCh38 pre-mRNA genome to generate gene by cell matrices for each library. Pegasus (https://github.com/lilab-bcb/pegasus), a python based program, was used to stringently filter cells, remove doublets by scrublet, integrate and batch-correct all libraries together. Initially we performed low-resolution analysis with top 10PCs and identified a low quality cell cluster with no canonical gene signature but high mito_percent and low ngenes. Then we re-ran the analysis with top 65PCs after removing these cells. Cells were removed if they expressed >6000 or <250 genes, or had >10% mitochondrially mapped reads. Dimensional reduction was done by PCA with n_genes and percent_mito were regressed out. Harmony (as part of the Pegasus suite) was used to integrate and batch correct libraries, Louvain clustering was performed to cluster the cells and visualize resulting clusters with UMAP.

### Cell type annotation

Cell types were annotated based on expression of known marker genes visualized via UMAP and by performing unbiased gene marker analysis (Table S1-3). Canonical genes were selected based on mouse and human studies as well as published reference atlas enriched genes (15–18). We integrated Velmeshev et. al. (13) dataset with our clusters through Seurat’s, R-based program (71) (https://satijalab.org/seurat/index.html), implementation of FindTransferAnchors and TransferData functions as detailed in primary manuscript as well as vignettes online (https://satijalab.org/seurat/articles/integration_mapping.html). In brief, the functions aim to build “anchors”, which are cell pairwise correspondences between single cells across the ‘reference’ datasets and ‘query’ datasets using PCA coordinates. Thus allowing integration of datasets into a shared space and serves as an effective matching tool of one cluster to another on the cell by cell bases (summarized in Table S1-2). We also used Azimuth, a web-based portal of the same Seurat functions above, from the Allen Brain Atlas (19) (https://azimuth.hubmapconsortium.org) to align our cell cluster to the human and mouse motor cortex clusters (BRAIN initiative annotation; *18*) (Table S1**-**2).

### Cortical Cell Composition Analysis

A cell type by subject count matrix of cell number per subject per cluster was generated from metadata, data was then transformed to cell proportions by total cells for each library, and log normalized. We then utilized the R-package LimmaVoom (72) to stringently ask if any clusters were significantly more or underrepresented in the ASD subjects in comparison to CTL subjects by implementing a linear model (Co-variables were included if a spearman correlation was above 0.25 against the top 5 clustering PCs): ∼diagnosis + Chemistry10X + age + PMI + totalReads. We repeated this analysis after removing individuals with known co-occurring Epilepsy to find very similar results (Table S3)

### Layer-Specific Spatial Transcriptomics Dataset (Nanostring Technologies)

We selected 4 CTL and 4 ASD subjects to embedded frozen cortex overnight in OCT (Tissue-Tek Cryo-OCT Optimum Cutting Temperature (OCT) #M-649078-1412) at −80C. We then equilibrated blocks to the −20C in a Leica cryotstat and collected 4 sections (at least 150uM apart from on another within the cortex) at 10uM thick sections onto Nanostring slides. Nanostring Technologies stained slides with Dapi (4′,6-diamidino-2-phenylindole) and GFAP (Glial fibrillary acidic protein) to help mark white matter boundaries from grey matter in Layer 6 and Layer 1/2. The sections were then treated with UV-photocleavable oligo-probes of about 18,000 genes from the Human genome. We assisted Nanostring technician, Jingjing Gong, to choose regions of interest or ROIs, we created 3-4 ROIs within grey matter for L1/2, L2/3, L3/4, L5/6 and three ROIs within White matter. The ROIs are then exposed to UV light which cleaves the probes releasing them for subsequent RNA sequencing. Following sequencing, the processing and normalization of data involves matching the raw probe counts to sequencing results to filter out unused probes and remove outliers then it is subjected to normalization to the third quartile.

#### Mapping gene activity signatures to layers

To look for spatial activity across the layer dataset using either cell-type annotation markers and or ASD DE genes: we first filtered each cell-type specific gene set or signature into small representative list to contain only positive unique genes to that set of comparisons. For cell-type specific ASD DE only significantly downregulated unique genes (FDR<0.05 and LOGFC above 0.2) were used to query layer activity for interneurons. For cell-type specific regulons, the top DE downregulated genes overlapping the regulon per neuronal type and the top DE upregulated regulon genes that are unique to MG2 were used to query layer count distribution. Linear mixed models were used to assess diagnosis effects and diagnosis × layer interaction effects using individual and nGene as random effect co-variables. *Post hoc* tests were assessed using estimated marginal means (least-squares means).

### Pseudobulk Expression Analysis by Cell Type

To correlate bulk tissue mRNA levels with nuclear RNA levels, we generated pseudobulk counts for each sample by adding counts from the same cell type. Then pseudobulk counts are normalized by variance stabilizing transformation method. To identify genes differentially expressed in ASD compared to control in each cell type, we examined covariates with top 5 PCs from normalized pseudo-bulk expression matrix. We identified the following covariates consistently correlated with top 5PCs for each cell type (age, PMI, BrainRegion, SeqBatch, Mito_perc and ngenes). We then randomly select subjects 500 times and calculated average beta to regress out effects of these covariates. Then we used limma-voom to identify differentially expressed genes for each cluster.

#### Single-cell RNAseq Bulkanized library

To correlate bulk tissue mRNA levels with nuclear RNA levels, we bulk-anized snRNA-seq data by aggregating all nuclear profiles by sample and compared bulk mRNA (7) FPKMs to normalized UMIs. We examined co-variates with top 5 PCs from normalized bulkanized matrix. We identified the following co-variates consistently correlated with top 5PCs for each cell type (age, PMI, SeqBatch, Mito_perc and ngenes) and calculated average beta to regress out effects of these covariates. Then we used limma-voom to identify differentially expressed genes ASD Vs CTL.

### Statistical tests

Unless otherwise stated all statistical test were run using R (4.1.0). Only P-values with post-hoc FDR or Bonferroni-corrected value of 0.05 were considered significant, unless otherwise indicated.

#### Module enrichment

We implemented a Fischer’s exact test to analyze the overlap of up regulated genes and down regulated genes for each cluster’s DE gene set in ASD. We filtered the DE gene lists to be FDR<0.05 and the kME of the module genes >0.5 to best capture the sn DE to module relationship. Odds ratio and estimated significance of overlap was determined using a background list of genes expressed in post-mortem brain.

#### Gene Ontology (GO)

We used PANTHER (Protein Analysis Through Evolutionary Relationships; http://pantherdb.org) to perform statistical overrepresentation test for DE from each cluster (Table S5-1) and for cell-type specific regulons (Table S8) utilizing fishers exact test with Bonferroni correction. We used brain-expressed genes as background. We only report GO terms that are significant (*corrected P = <0.05*). All GO ontologies categories (cellular composition, biological process, molecular function, REACTOME pathways, protein function) are reported for each cell and all were used to interpret DE changes.

We also implement SYNGO (https://www.syngoportal.org) under the same settings as above to verify synaptic enrichments in neurons as well as validate pre-and post-synaptic gene functions (Table S5-2,3).

#### Protein-Protein Interaction networks

DAPPLE (Disease Association Protein-Protein Link Evaluator; *73*) results for evaluating the significance of PPI networks of the cell-type specific regulons were all done using 1,000 permutations (within DAPPLE parameter) and *P values* < 0.05 were considered significant.

#### Risk Gene overlap in bulk DE

To estimate overlap of ASD genetic risk factors with DE we used the hypergeometric test, we used the list of DE in scBulkanized dataset and list of all genes expressed. These lists were overlapped with all genes in the SFARI Gene Module database having evidence of type 1, type 2, or syndromic genetic association with ASD (https://gene.sfari.org/database/gene-scoring).

#### Risk Gene overlap in regulons

We used Fisher’s exact tests to estimate the significance of overlap between genes within select regulons and significantly differentially expressed (DE, FDR<0.05) genes by cell-type, split by upregulated (log2FC>0.2) or downregulated (log2FC<-0.2) using a background list of genes expressed in postmortem brain (expressed in at least 100 cells, 15,370 genes). Estimated p-values were FDR corrected. Similarly, we used Fisher’s exact tests to compare the genes associated with Intellectual Disability (57), Fragile × Mental Retardation Protein membership (58), or a SFARI genes either considered levels 1, 2, or Syndromic with TFs/drivers of significant regulons within each cell-type. Odds ratio and estimated significance of overlap was determined using a background list of genes expressed in post-mortem brain.

#### Common Variation enrichment in Regulons

Partitioned heritability of regulons and ASD PRS (55) was assessed using LD Score Regression v1.0.0 (74). Heritability was calculated by comparing the association statistics for common genetic variants falling within regulatory elements associated with specific regulon genes, with the LD-score, a measure of the extent of the LD block. First, an annotation file was created, which marked all SNPs that fell within the regulatory elements for each regulon. LD-scores were calculated for these SNPs within 1 cM windows using the 1000 Genomes EUR data. These LD-scores were included simultaneously with the baseline distributed annotation file from Finucane et al., 2015 (74). The enrichment coeffiencient was calculated as the heritability explained for ASD common variation within a given annotation divided by the proportion of SNPs in the genome and subject to Bonferonni correction.

### SCENIC Regulatory networks

We employed PySCENIC (*47*; https://pyscenic.readthedocs.io) on a subset of cells within our dataset (10K cells, randomly subsetted). Prior to subsetting, the dataset was filtered to remove cells with fewer than 200 genes as well as to remove genes expressed in fewer than 100 cells. We then constructed gene regulatory networks (GRNs) informed by a list of all Transcription Factors (RCIStarget). GRNs are then pruned to only include downstream targets that have a TF-binding motif within 10KB of the TSS. Finally, area under the curve scores (AUC) were scored across 100K cells and z-scored AUC scores were calculated. Regulons wereconsidered expressed in a cell-type if they had an average z-score >0. AUC z-score values of expressed regulons within each cell-type were compared between ASD and CTL using t-tests and FDR corrected p-values (corrected within cell-type based on the number of expressed regulons).

#### Regulon overlap of transcriptional changes in each cell type

The list of TF-drivers of regulons that were significantly DE in at least one cell type in ASD (217 regulons out of total 337 regulons) were overlapped with each cell types’ ASD DE genes to identify which TF’s expression in down or up regulated. These TF-lead regulons were selected to calculate the percentage of overlap between the total regulon target genes and DE genes of a given cell type. To consider a ‘top’ regulon overlap we investigated: 1) the percent the regulon overlaps with highly upregulated genes and/or highly downregulated genes within that cell type (FDR<0.05;LOGFC>0.2;LOGFC<-0.2) and 2) the significance/OR from a fisher exact test (above) to all up-or all downregulated significant DE genes (FDR<0.05) across all cell types. Using this criteria we focused on ‘top’ regulons with the most significant and highest percent overlap of DE genes for a given cell type.

### ScATAC sequencing and data analysis

Nuclei were extracted as above simultaneously with 22 samples subjected to scRNAseq – consisting of 12 ASD samples and 10 CTL individuals that were balanced by age, PMI and cause of death (See Table S1-6 for scATACseq meta data). Libraries were aligned to reference genome with 10X Genomics software, Cell Ranger (cellranger-atac count; https://support.10xgenomics.com/single-cell-atac/software) then CTL libraries and ASD libraries were aligned together using the cellranger-atac aggr function. We utilized Signac (*75*; https://stuartlab.org/signac/), R-based program by the same lab as Seurat, to stringently filter out low quality cells through nucleosome banding score < 4 and TSS < 2 leaving 41,683 ASD - and 31,754 CTL high quality cells from each dataset. We merged ASD and CTL objects together by union of peaks from both and then re-called peaks so the same set of features are accounted across both datasets. Now merged, we normalized with term frequency inverse document frequency (TF-IDF), reduced dimensionality using partial singular value decomposition, and clustering omitting LSI1 to generate 30 clusters. To annotate the clusters, we first created a gene activity score matrix where gene ‘expression’ levels are roughly computed from fragment counts within gene body elements (See: https://stuartlab.org/signac/articles/). We then used the snATACseq ‘gene activity matrix’ to integrate with our fully annotated snRNAseq clusters through implementing Seurat’s ‘FindTransferAnchors’ function as described above. We find 30 clusters in the scATACseq dataset that fit the profile of the snRNAseq clusters, given that the scATACseq data is ∼6X smaller than snRNAseq it is expected to not have as many clusters, specifically they do not have cluster OPCS_3, ASTRO_2, and a number of small L56 excitatory clusters. These cluster likely did not reach enough number in the ATAC dataset to form a separate cluster. We then used spearman correlation to assess how closely related the cluster-specific marker gene profiles from the snATACseq clusters (from ‘gene activity matrix’) with our snRNAseq cluster marker gene profiles.

### Peak calling and prediction of TF activity in snATAC-seq data

We merged reads from individual cells of the same snATAC clusters per sample (https://github.com/timoast/sinto) with the following commands: sinto filterbarcodes --bam {bam} --cells {barcode_path} --outdir {subset_dir} --nproc 4. Pseudobulk replicates were further created with samtools merge by disease condition. Peak calling was performed using MACS2 (https://github.com/macs3-project/MACS) in Signac, resulting in a reproducible, non-overlapping peakset. To infer transcription factor activity, we performed TF foot-printing analysis on the consensus peak-set using TOBIAS (53), as described in our previous study (54). Briefly, this method starts with Tn5 bias correction using the TOBIAS ATACorrect module, subtracting the background Tn5 insertion cuts highlighting the effect of protein binding. The foot-print score was calculated by TOBIAS ScoreBigWig, which measures both accessibility and depth of the local foot-print, thus correlating with the presence of a TF at its target loci, and the chromatin accessibility of the regions where this TF binds. To match foot-prints to potential TF binding sites, and to estimate TF binding activity on its target loci, we applied TOBIAS BINDetect module to the corrected ATACseq signals within peaks, with TF motif PWMs from TRASFAC Pro. Many transcription factors are represented by more than one motif. To avoid motif redundancy, we clustered the motifs based on their sequence similarity using TOBIAS ClusterMotifs, and chose one motif per TF that is most similar to others in the same cluster TOBIAS BINDetect compares the positions and activities of TF foot-printed sites in disease or control per clusters. Each foot-print site was assigned a log2FC (fold change) between two conditions, representing whether the binding site has larger/smaller TF foot-print scores in comparison. To calculate statistics, a background distribution of foot-print scores is built by randomly subsetting peak regions at ∼200bp intervals, and these scores were used to calculate a distribution of background log2FCs for each comparison of two conditions. The global distribution of log2FC’s per TF was compared to the background distributions to calculate a differential TF binding score, which represents differential TF activity between two conditions. A *P-value* is calculated by subsampling 100 log2FCs from the background and calculating the significance of the observed change. By comparing the observed log2FC distribution to the background log2FC, the effects of any global differences due to sequencing depth, noise etc. are controlled. To visualize, we used soGGI to plot TF foot-prints, and bar graphs to show the global TF foot-print activity changes comparing ASD versus Control.

## Acknowledgements

Funding: The work is funded by the U.S. National Institute of Mental Health (NIMH) (grants R01-MH109912, D.H.G.;R01-MH100027, D.H.G.; U01-MH115746, D.H.G.; R01-MH116489, D.H.G.; R01-MH100027, P50HD103557 D.H.G.; R01-MH110920, the Simons Foundation for Autism Research Initiative (SFARI grant 675474, D.H.G.); B.W. was partially supported by N.I.H. postdoctoral training T32NS048004 program in Neurobehavioral Genetics at UCLA. Data were generated as part of the PsychENCODE Consortium (U01MH116489 awarded to M. Gerstein (Yale), D. Geschwind (UCLA), and K. White (Yong Loo Lin School of Medicine, NUS)).

Also, we thank program and support staff, in particular L. Bingaman, D. Panchision, A. Arguello, and G. Senthil, for providing institutional support and guidance for this project. We also thank Michael Gandal, Mark Gerstein, Kevin White, Sofia Gaynor, Prashant Emani, and members of the Geschwind lab for helpful discussions. Cell figures were created originally by B.W. with BioRender.com. Brain tissue for the study was obtained from the following brain bank collections: University of Miami Brain Endowment Bank, University of Pittsburgh Brain Tissue Donation Program, University of Maryland Brain and Tissue Bank and others under the NIH NeuroBioBank and the Autism Brain Net (ABN) project supported by the Simons Foundation for Autism Research Initiative (SFARI; now containing the previous Harvard Brain Bank Tissue Resource Center as part of the former Autism Tissue Project (ATP)). We also thank Wes Goldman and Jingjing Gong at Nanostring Technologies and their colleagues for assistance with conducting the spatial transcriptomics and guidance in analyzing the layer-specific transcriptomic data.

## Author Contributions

Data were generated by B.W., D.Q., S.M., J.L., L.B., N.H. and D.H.G. and the PsychENCODE Consortium, Data analysis was performed by B.W., L.B., J.G., Y.C, R.K., and D.H.G. The manuscript was written by B.W., L.B., Y.C., and D.H.G. The work was supervised by D.H.G.

## Data availability

The source data described in this manuscript are available via the PsychENCODE Knowledge Portal (https://psychencode.synapse.org/). The PsychENCODE Knowledge Portal is a platform for accessing data, analyses, and tools generated through grants funded by the National Institute of Mental Health (NIMH) PsychENCODE Consortium. Data is available for general research use according to the following requirements for data access and data attribution: (https://psychencode.synapse.org/DataAccess). For access to content described in this manuscript see: https://www.synapse.org/#!Synapse:syn51032009/datasets/.

## List of Supplement materials

**<u>Figs. S1 to S9</u>** Supplementary Figures pg. 2-19

**<u>Tables S1, S2, S3, S4, S5, and S6</u>** Supplementary Table contents pg.20-21

**Figure S1.**
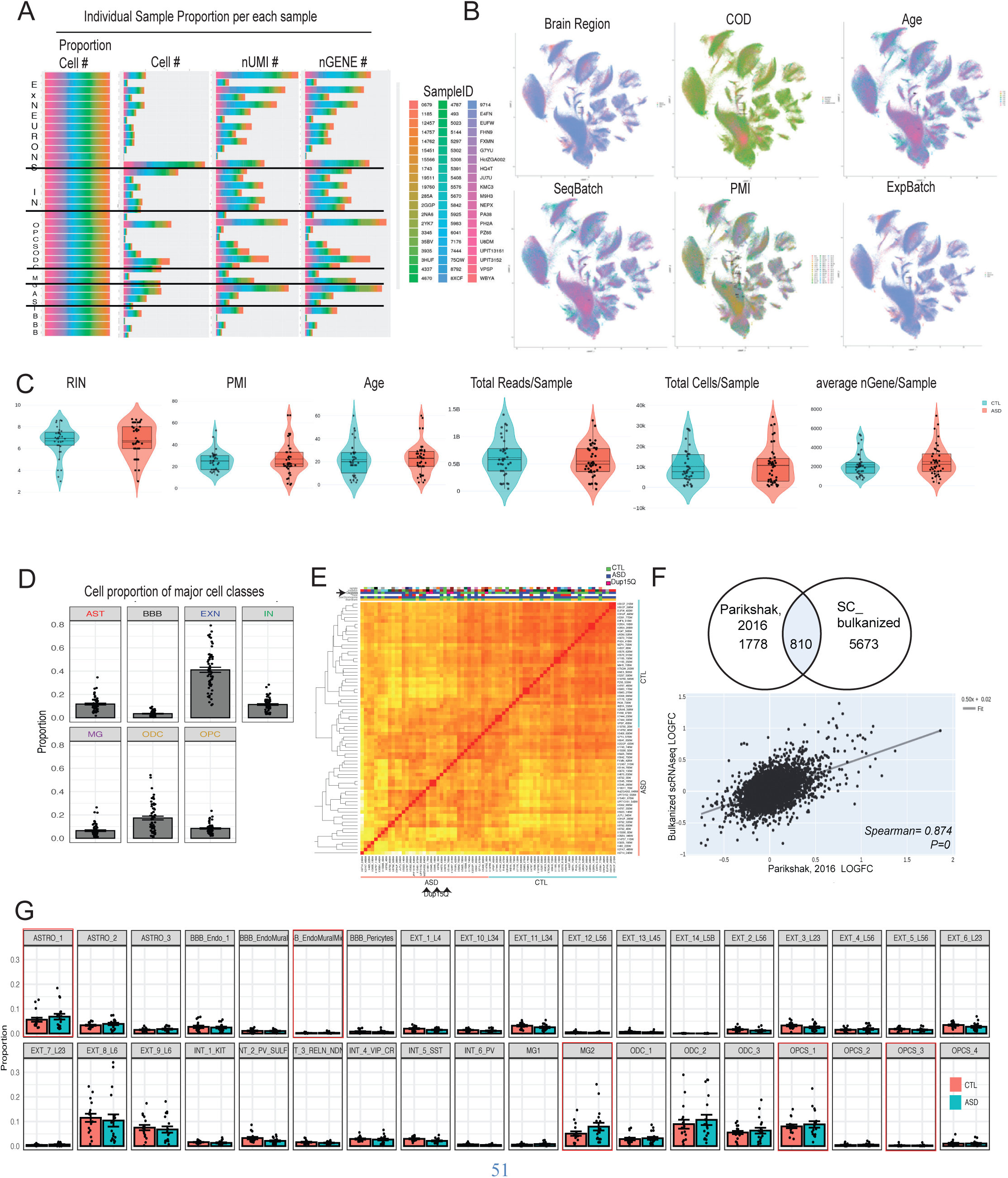
Meta data and cell composition of dataset. A) Individual cell proportion: per cell cluster (left), per raw cell number per cluster (middle left), per nUMI/total RNA (middle right), and per nGENE per cluster (right). B) Individual UMAPs of cells colored by meta data variable: Brain region (top left), COD (top middle), Age (top right), SeqBatch (bottom left), PMI (bottom middle), and ExpBatch (bottom right). C) Violin plots of meta data values for CTL (blue) and ASD (red) samples from left to right: RIN, PMI, Age, total reads per sample, and average nGENE per sample. D) cell cluster proportions of major cell classes, E) Heatmap of top variable genes illustrating that DUP15Q and ASD samples cluster together on the left side separately from most CTL samples on the right. F) Plot of current study DE genes LOGFC against previous bulk DE genes LOGFC (6) at FDR<0.1 of 810 shared genes (bottom) spearman correlation of 0.87 (*P = 0*). G). Cell proportions for each subtype cluster from each sample. *Abbreviations: nUMI – number of unique molecular identifier, nGene – number of genes, COD – cause of death, SeqBatch – sequencing batch, ExpBatch-experimental batch, PMI – post-mortem interval, CTL – control individuals, ASD – autism spectrum condition, RIN – RNA integrity number, DUP15Q – individuals with duplication of 15Q and ASD, DE – differential expression, LOGFC – log transformed fold change ASD versus CTL*.

**Figure S2.**
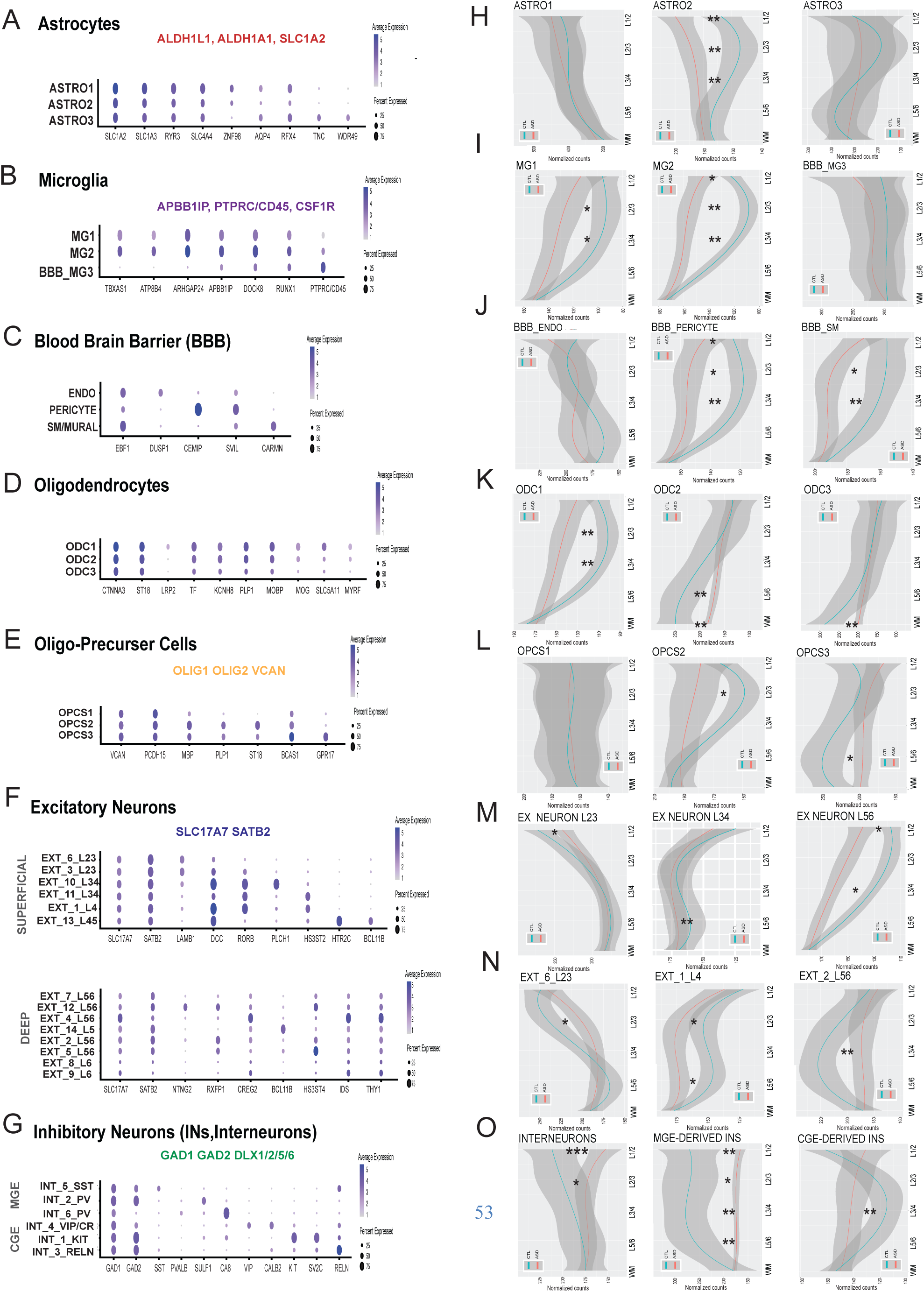
Cluster specific marker gene expression predicts spatial distribution in cortical layers. A-G) Dot plot of marker genes for each cluster where the size of the circle depicts the percentage of cells within the cluster that express the gene and the blue color depicts the level of expression. Shared genes are listed in bold above the dot plot. H-O) Spatial distribution across cortical layers of cell cluster-specific list of marker genes unique to that cell with normalized counts from the CTL brain (blue line) and ASD brain (red line). *0.01<*P*<0.05 **0.001<*P*<0.01 \*\*\**P*<0.001.

**Figure S3.**
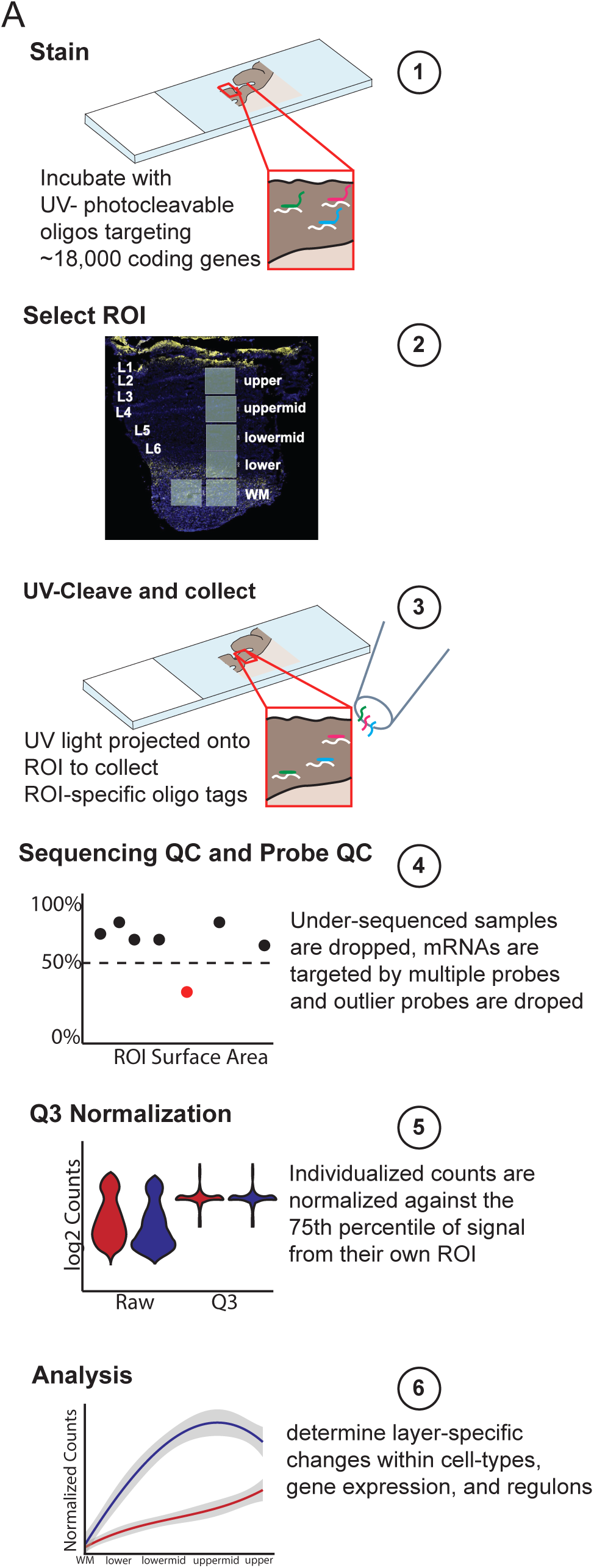
Nanostring protocol for spatially selective transcriptomics. A) Postmortem tissue from ASD and CTL individuals (n=4 subjects each, 4 sections per cortex) was stained for ∼18,000 protein coding genes with UV-photocleavable oligo tags, DAPI and GFAP antibodies (1). Regions of interest (ROIs) were drawn based on laminar structure of the cortex (2). Oligos within a specific region are cleaved, collected, and sequenced (3). Sequenced reads are matched to target genes, and samples with less than 50% sequencing saturation are dropped. mRNAs are targeted by multiple probes and any outlier probes for a given target are dropped from further analysis (4). Counts are normalized to the 75th percentile of all counts for that ROI (5) and samples are analyzed using linear mixed models to determine ASD vs. CTL changes across layers of the cortex (6).

**Figure S4.**
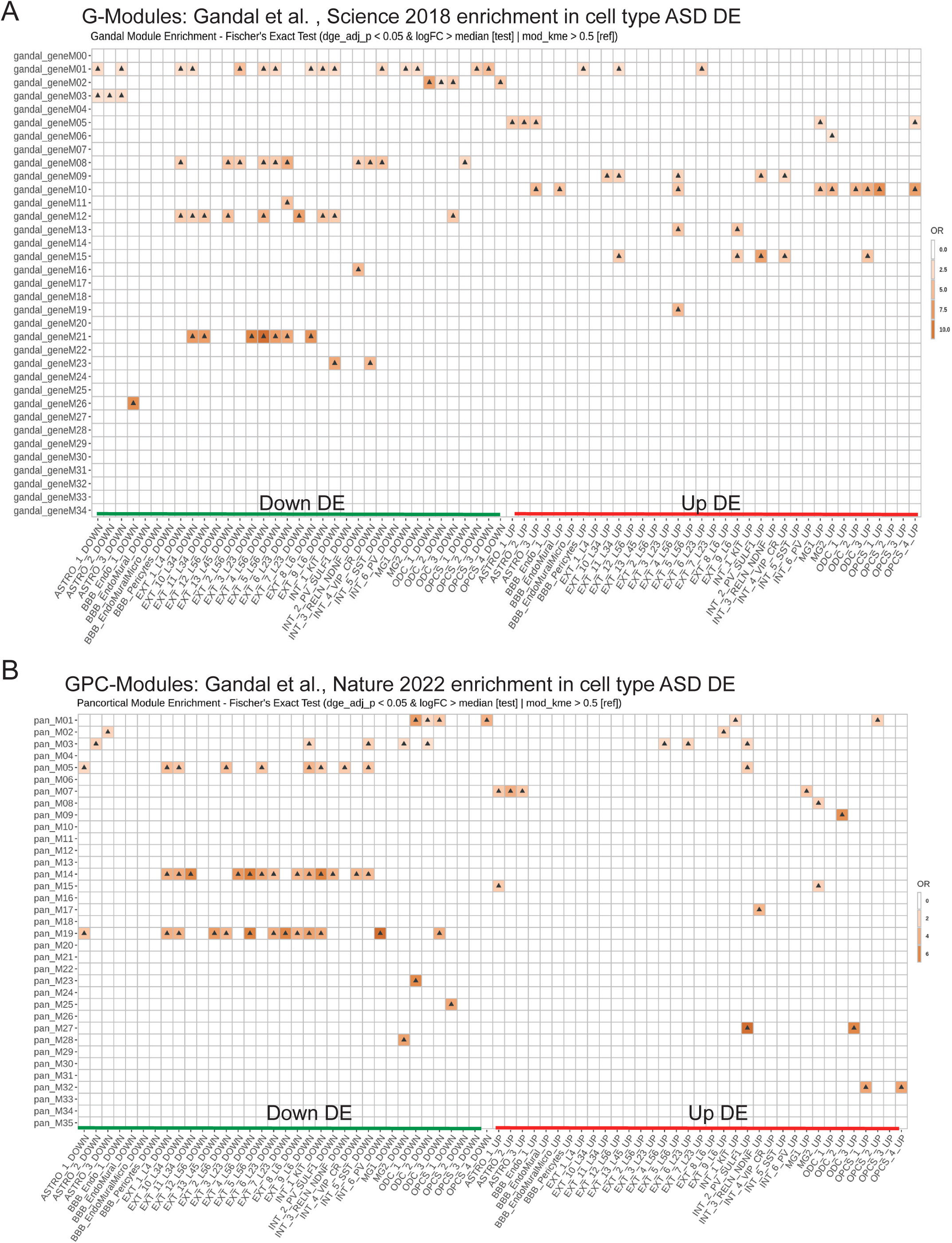
Cell-type specific co-expression module associations from ASD bulk studies. A-B) Heatmap of OR enrichment of (A) ‘G-modules’ (from Gandal et. al, 2018 (8)) or (B) ‘GPC modules’ (from Gandal et. al., 2022 (14)) and cell-type specific upregulated DE genes in red and downregulated DE genes in green. Only significant association are shown as indicated by black triangle. *Abbreviations: G – Gandal, GPC – Gandal pancortical, DE – differential expressed*.

**Figure S5.**
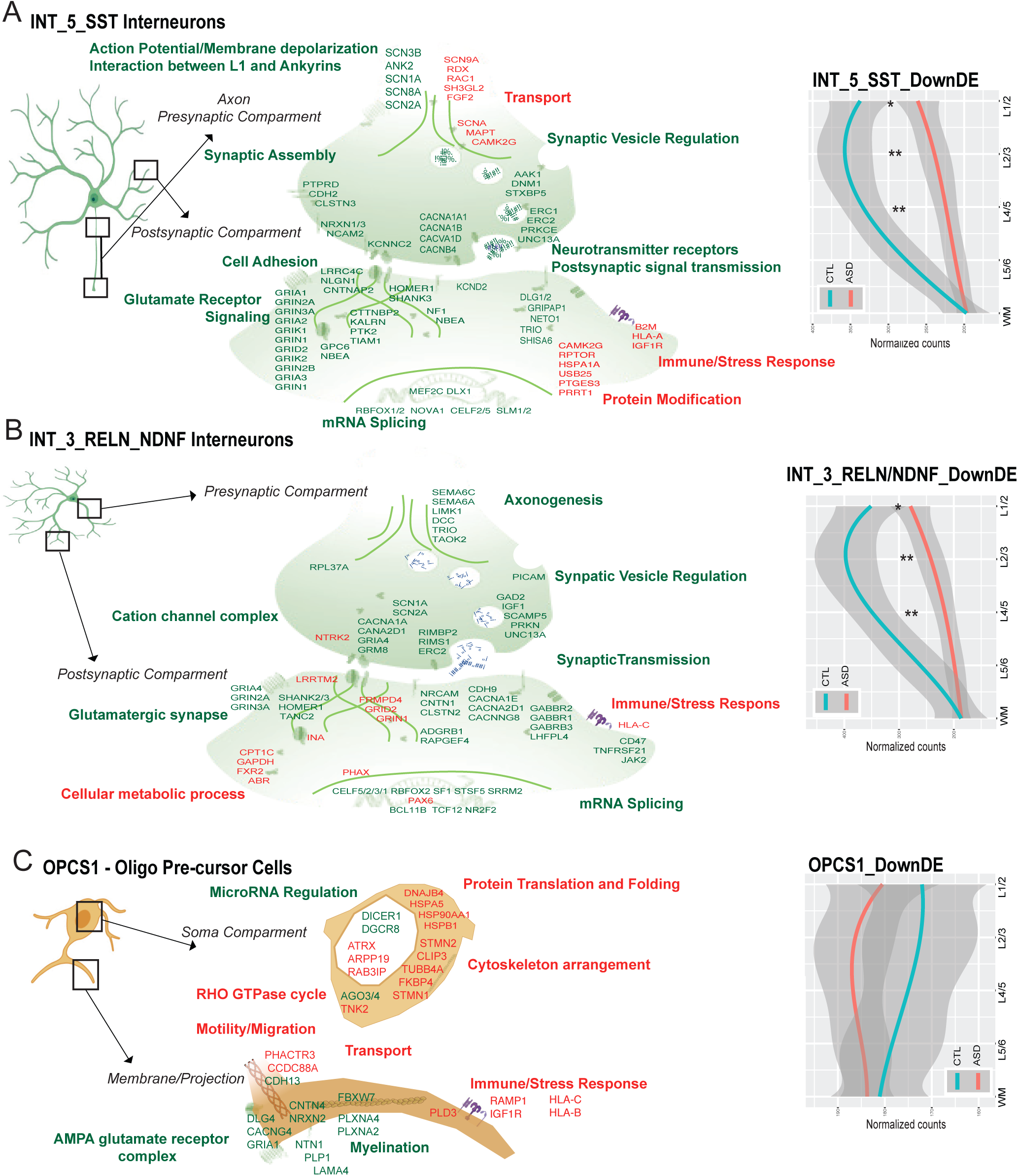
Cell-type specific transcriptional changes in INs and OPCs. A-B) Schematic of SST+ neurons (A) or CGE-derived *RELN*+ neurons (B) in green, representing the cell soma, dendrites, and axonal projection; black square indicates putative location enlarged example of the presynaptic axonal compartment on top and enlarged example of postsynaptic dendritic compartment on bottom. The most significant up regulated GO terms and pathways (red) and their accompanying DE genes (red) are depicted; down regulated GO term and pathways (green) and their accompanying DE genes (green). The panel on the right depicts the spatial distribution across cortical layers of cell cluster-specific down regulated DE genes unique to that cluster with normalized counts from the CTL brain (blue line) and ASD brain (red line) with Layer (L) 1/2 on top followed by L2/3, L4/5, L5/6 and WM on bottom. C) Cartoon of OPCS, OPCS1, with a black box indicating putative location of the DE genes in enlarged myelinating membrane/ projection, top, and soma compartment, bottom. The most significant up regulated GO terms and pathways are in red and their accompanying DE gene examples are labeled in red, down regulated GO term and pathways are marked in green and their accompanying DE gene examples are labeled in green. The panel on the right depicts the spatial distribution across cortical layers of cell cluster-specific down regulated DE genes unique to that cluster with normalized counts from the CTL (blue line) and ASD (red line) brain with Layer 1/2 on top followed by L2/3, L4/5, L5/6 and WM on bottom. *0.01<*P*<0.05 **0.001<*P*<0.01 \*\*\**P*<0.001. *Abbreviations: CGE – caudal ganglionic eminence, GO – gene ontology, DE – differentially expressed, WM – white matter, OPCS – oligo-precursor cells, ODC – oligodendrocytes, CTL – control individuals, ASD – autism spectrum condition*.

**Figure S6.**
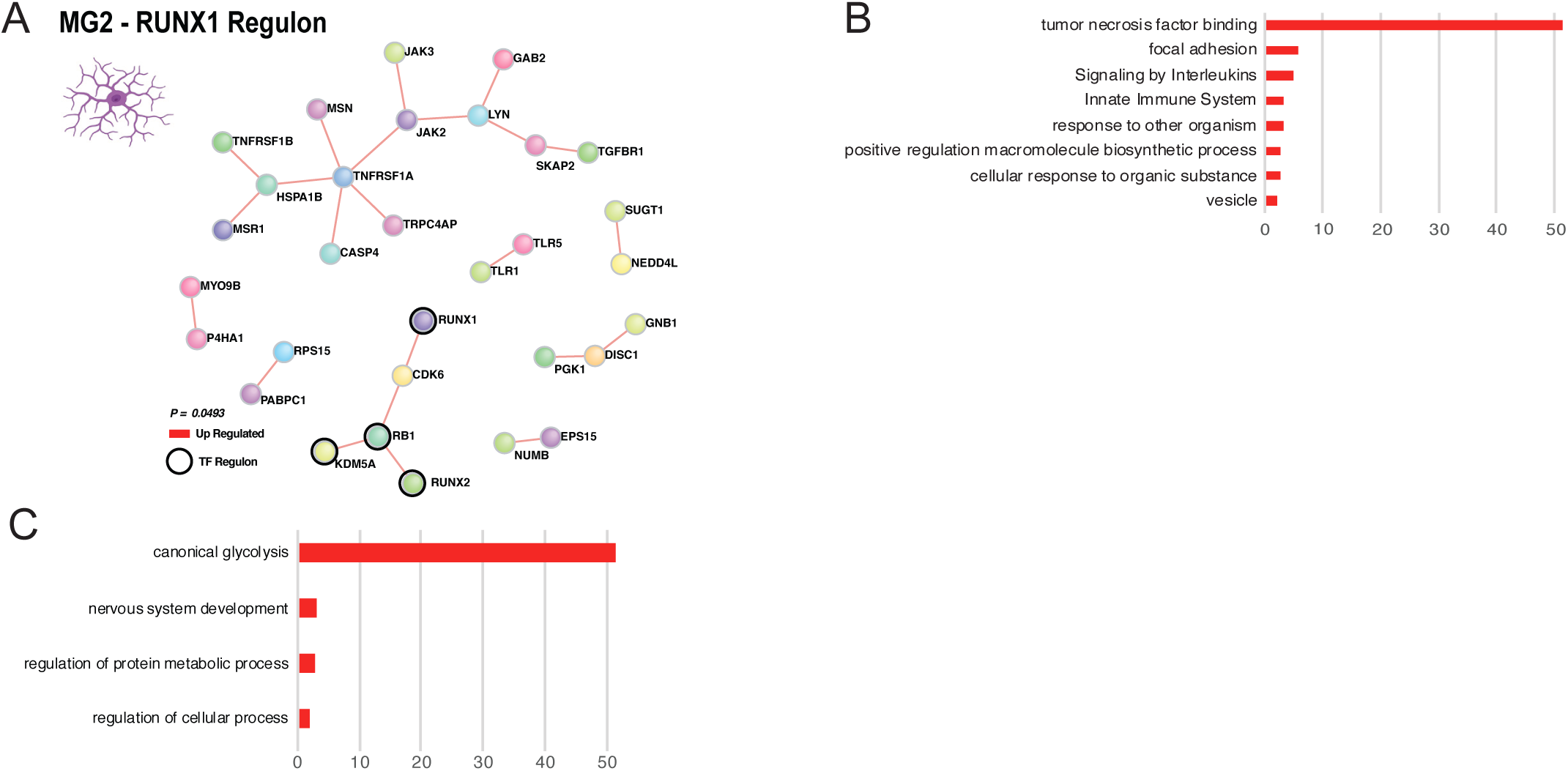
MG2 top regulons account for substantial portion of upregulated genes related to inflammatory signaling. A) PPI network or *RUNX1* upregulated genes within MG2 cells includes genes related to reactive inflammatory state and other TF – regulon drivers (black cirled) B) Significant GO terms associated with upregulated DE genes encompassing the *RUNX1* regulon in MG2 cells. C) Significant GO terms associated with upregulated DE genes encompassing the *ETV6* regulon in MG2 cells. *Abbreviations: MG – microglia, DE – differentially expressed, GO – gene ontology*.

**Figure S7.**
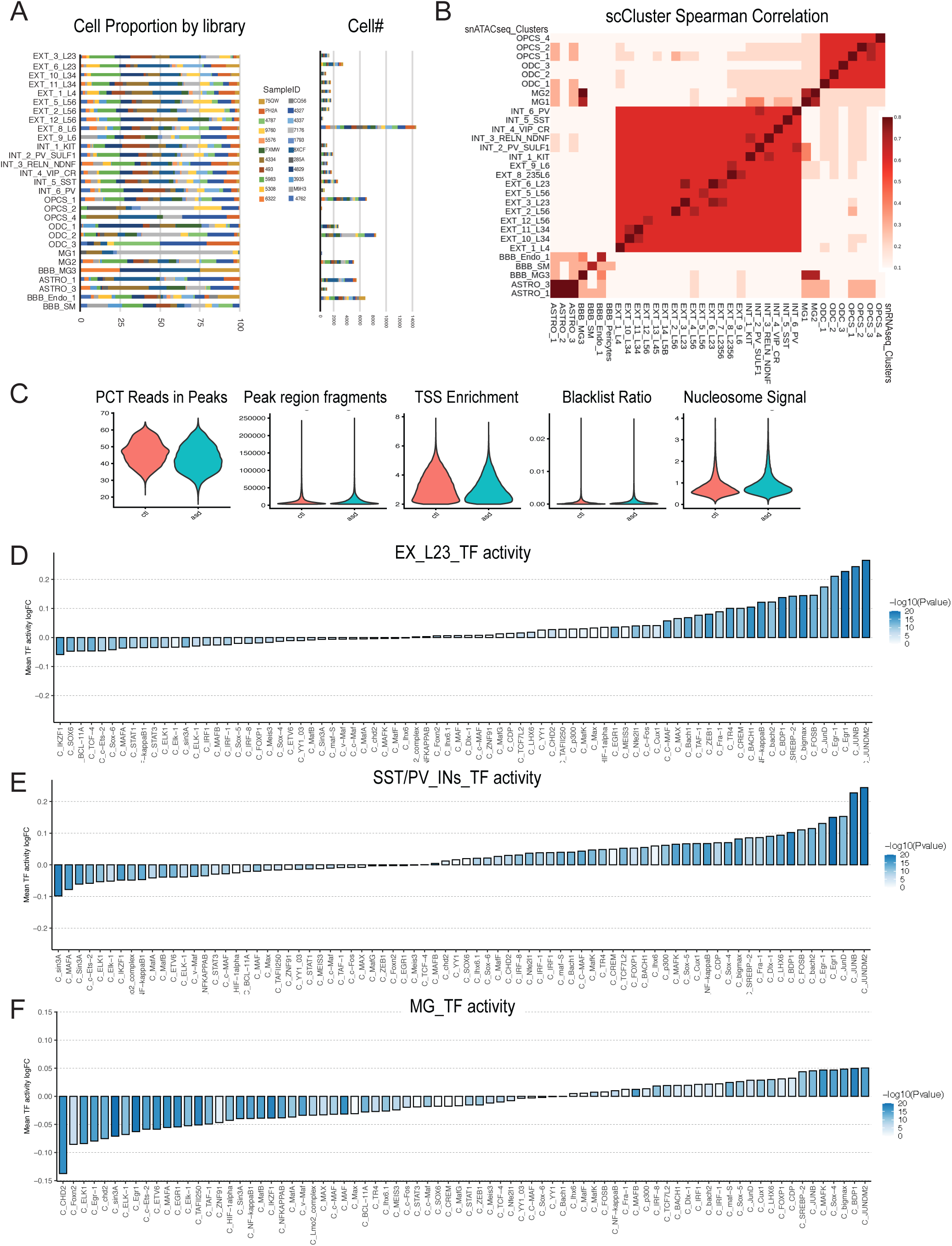
ScATACseq dataset composition and TF footprints that are significantly altered by cell type. A) Individual sample proportion per cluster in the scATACseq dataset. B) High correlation between scATACseq cluster profiles and scRNAseq cluster profiles (Spearman correlations (red), C) Violin plots of quality metrics across CTL cells (blue) and ASD cells (red) of scATACseq dataset. D-F) Bar plot of fold change of significantly altered TF footprints shown to regulate DE genes in aggregate (D) Ex Neurons, (E) *SST/PV* INs, and (F) MG cell types. The height of the bar indicates the level of fold change and significance indicated (blue). *Abbreviations: CTL – control individuals, ASD – autism spectrum condition, TF – transcription factor, Ex Neurons – excitatory neurons, INs-interneurons, DE – differential expression*

**Figure S8.**
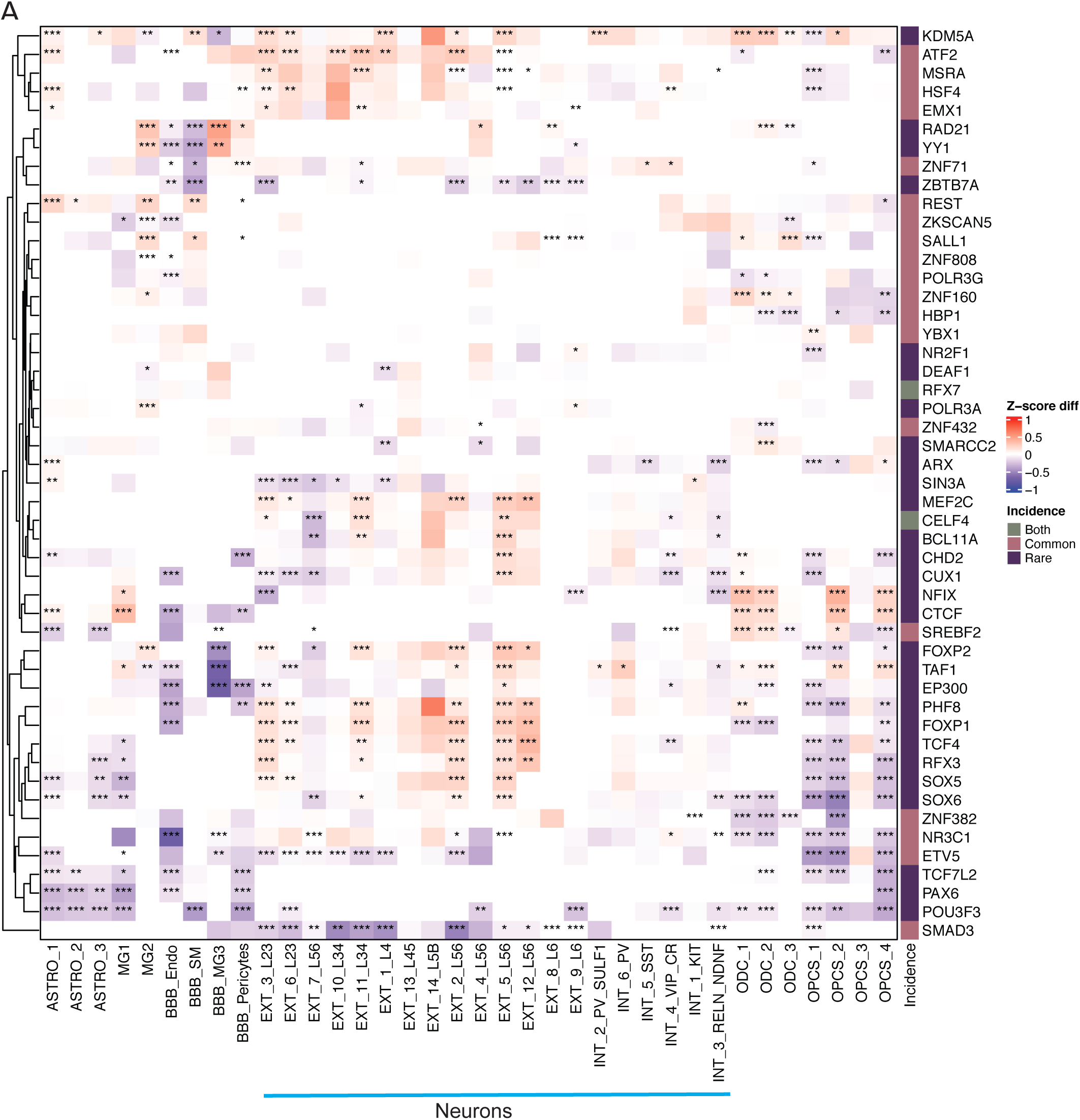
ASD risk gene - and PRS – enriched regulon networks significantly DE in ASD cell types. A) Heatmap of significantly altered regulon expression in ASD by cell type showing Z score diff of AUC values where red indicates increase activity of network genes and blue indicates decreased expression of network genes within each cell type. Regulons are marked whether they are headed by TFs that are rare risk genes for ASD (SFARI genes) in purple or enriched for ASD PRS (common variation from Grove et al., 2019, *55*) in pink or both in grey. Neuronal clusters are labeled by blue bar. *0.01<*P*<0.05 **0.001<*P*<0.01 \*\*\**P*<0.001. *Abbreviations: SFARI – Simons Foundation for Autism Research Initiative*.

**Figure S9.**
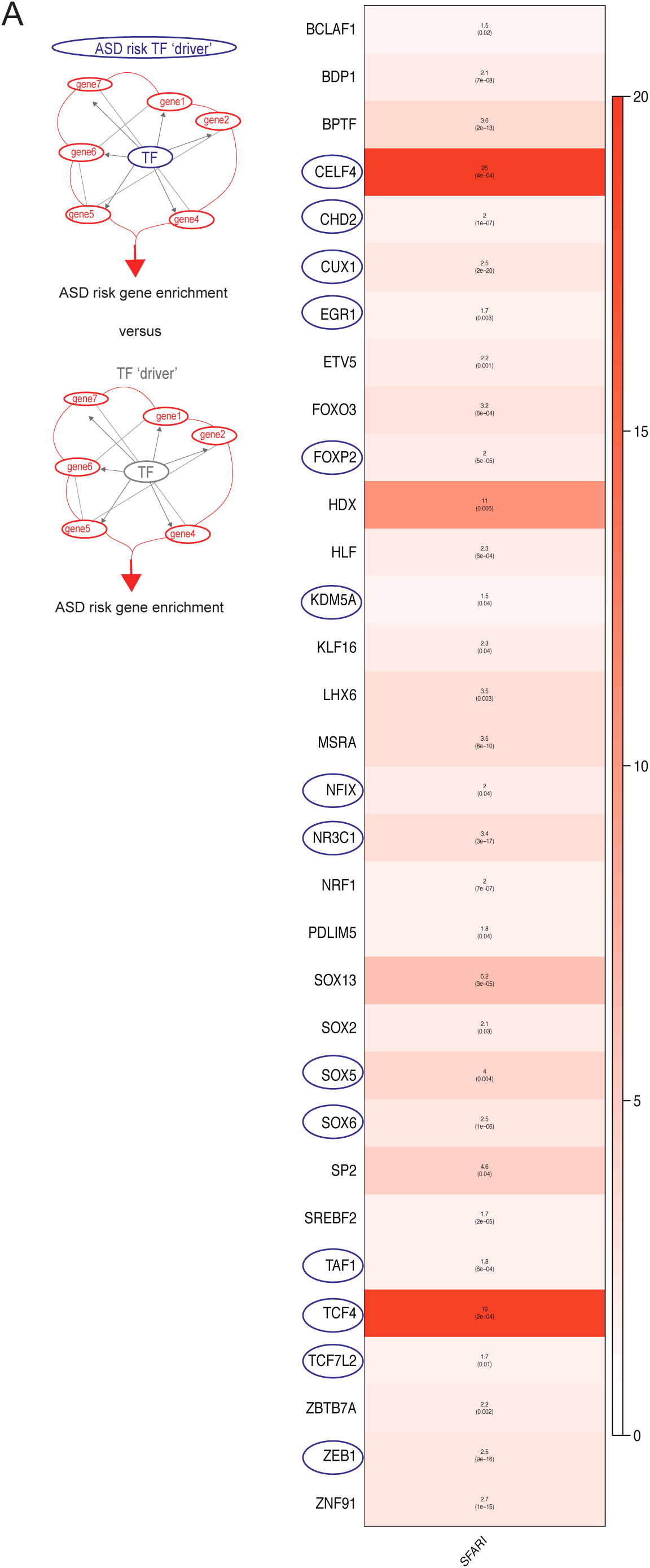
Regulons driven by ASD risk TF drivers are enriched for ASD risk. A) Heatmap of significant ASD rare risk gene enrichment (OR; red) from all regulons, OR and *P-value* included (black). *Abbreviations: ASD – autism spectrum condition, TF – transcription factor, OR – odds ratio*.

**Table S1:** Tab 1) scRNA Meta data by sample; 2) Cluster Annotation; 3) Cluster marker genes; 4) Velmeshev et. al. 2019 (13) comparison; 5) scATAC Meta data by sample (892 KB)); Link to file: https://www.dropbox.com/scl/fi/hzt672gjcp8v73gkyes1s/Table_S1_Wamsley_PEC_2_21_23.xlsx?dl=0&rlkey=o7worwpmfhj2b7rcmpsx0fcxq

1. scRNAseq Meta data by sample: biological and technical properties by individual samples; 2) Cluster cell type annotations equivalents from Allen brain atlas (19), Brain Initiative (18), and Velmeshev et al., 2019 (13); 3) Cluster specific marker genes, 4) Comparison of study components to Velmeshev et al., 2019 (13); 5) scATACseq Meta data by sample: biological and technical properties by individual samples

**Table S2:** Meta data per cell: Cell barcodes and meta data biological and technical properties by cell (101.2 MB); Link to file: https://www.dropbox.com/scl/fi/g85bursqyf5ycr1lndacb/Table_S2_Wamsley_PEC_2_21_23.xlsx?dl=0&rlkey=596xacoe1w890gy6jttxqfxfr

**Table S3:** Tab 1) Cell composition LV w Correction; 2) Cell composition LV w/o Correction (131 KB)); Link to file: https://www.dropbox.com/scl/fi/sufxn0g3qso7lfm5qnw0c/Table_S3_Wamsley_PEC_2_21_23.xlsx?dl=0&rlkey=9y7geir8fsmqau0m9sted0b2z

1. Cell composition analysis results with LimmaVoom (LV) including co-variables; 2) Cell Composition analysis result with LimmaVoom (LV) without corrections including all samples

**Table S4:** Tab 1) All module enrichments in cluster ASD DE; 2) Gandal 2018 G-Modules genes; 3) G-Modules GO; 4) Gandal 2022 G-PC Modules and; 5) GPC Modules GO analysis results (21.7 MB); Link to file: https://www.dropbox.com/scl/fi/9gfn79s9nehprpyah3a1z/Table_S4_Wamsley_PEC_2_21_23.xlsx?dl=0&rlkey=ykz7rm2pl1dcrxow0e6oetjmr

1. Fisher exact results for Module enrichment in ASD DE genes in each cell cluster; 2) Gandal et al., 2018 (8) gene modules and kME bulkRNAseq values; 3) Gandal et al., 2018 (8) module Gene Ontology (GO); 4) Gandal et. al., 2022 (14) gene modules and kME bulkRNAseq values; 5) Gandal et al., 2022 (14) module Gene Ontology (GO).

**Table S5:** Tab 1) Cell type specific GO; 2)SYNGO-BP; 3)SYNGO-CC (49 MB); Link to file: https://www.dropbox.com/scl/fi/hmzzwzsr3jxds8bb3gxeu/Table_S5_Wamsley_PEC_2_21_23.xlsx?dl=0&rlkey=fry4r9fg1q1m924u7tbvf4u31

1. Significant Gene Ontology for all cell types ASD DE; 2) SYNGO – BP, biological process, annotations for neurons; 3) SYNGO-CC, cellular component, annotations for neurons

**Table S6A:** Tab 1-19) Neuronal cluster specific ASDvCTL DE genes (27.8 MB): Link to file: https://www.dropbox.com/scl/fi/d90tusc7qucwzy1vnp0ry/Table_S6A_Neuronalclusters_DE_genes_2_21_23.xlsx?dl=0&rlkey=otokcg1mktzscer6wbjr2wknk

1-19) Differential Gene expression per neuronal cell type

**Table S6B:** Tab 1-16) Glia cluster specific ASDvCTL DE genes (21.8 MB); Link to file: https://www.dropbox.com/scl/fi/x8megov2qb05pmlshhemd/Table_S6B_Gliaclusters_DE_genes_2_21_23.xlsx?dl=0&rlkey=wqjdrz2o0u3faeo0edbt1nys5

1-16) Differential Gene expression per glia cell type

**Table S7: ’**Bulkanized’ ASD DE genes and stats from scRNAseq data (4.6 MB); Link to file: https://www.dropbox.com/scl/fi/ezrtfwh1g7vti4zet4l68/Table_S7_Wamlsey_2_21_23.xlsx?dl=0&rlkey=yzggtbq229zo9djowcmpfypqk

**Table S8:** Tab 1) All Regulons, 2) Binary matrix of Regulons per cell type, 4) LDSC with ASD GWAS (55) results per Regulon, 5) Cell-type specific regulons in INT_5_SST, 6) Cell-type specific regulons in EXT_6_L23, 7) Cell-type specific regulons in MG2 (657 KB); Link to file: https://www.dropbox.com/scl/fi/f1yurancjw3qz5e68li93/Table_S8_Wamsley_2_21_23.xlsx?dl=0&rlkey=41nvp0zi1qj2y63o10y5pesri

1. Regulons listed as columns; 2) Binary matrix of significant differential Regulons per cell (1-significantly changed in ASD, 0-non-significant or no activity change); 3) AUC scores ASD vs CTL per cell; 4) LDSC coefficient score and pvalues per regulon with ASD GWAS (55); 5) Regulons overlapping ASD DE within INT_5_SST; 6) Regulons overlapping ASD DE within EXT_6_L23; 7) Regulons overlapping ASD DE within MG2

**Table S9:** AUC scores ASD v CTL cells (465.1 MB); Link to file: https://www.dropbox.com/scl/fi/hz3mpkq16k61zfzciwbau/Table_S9_Wamsley_PEC_2_21_23.xlsx?dl=0&rlkey=q3ou0vznqkcir4wexdysu7wfe

**Table S10:** 1) TF foot-printing annotation for aggregated Ex Neurons (56.4 MB); Link to file: https://www.dropbox.com/scl/fi/s7lnsg0xe1uc3jtl8fzze/Table_S10_Wamsley_2_21_23.xlsx?dl=0&rlkey=qc5qz3cmp9z5ynpa3jy1cqmvi

1. TF footprint annotation and metrics for changing footprints of Regulon drivers regulating DE genes in Ex Neuron

**Table S11:** TF foot-printing annotation for aggregated MG cell types (57 MB); Link to file: https://www.dropbox.com/scl/fi/nk0eywpk7f9cjjiw5ntp4/Table_S11_Wamsley_PEC_2_21_23.xlsx?dl=0&rlkey=ha26dgy08w0wjtortfquartwf

1. TF footprint annotation and metrics changing footprints of Regulon drivers regulating DE genes in MG

## References

1. D. Wang et al., Comprehensive functional genomic resource and integrative model for the human brain. Science. 362, 8464–15 (2018).

2. M. Li et al., Integrative functional genomic analysis of human brain development and neuropsychiatric risks. Science. 362, 7615– (2018).

3. N. N. Parikshak et al., Integrative Functional Genomic Analyses Implicate Specific Molecular Pathways and Circuits in Autism. Cell. 155, 1008–1021 (2013).

4. P. Rajarajan et al., Neuron-specific signatures in the chromosomal connectome associated with schizophrenia risk. Science. 362, 4311– (2018).

5. M. J. Gandal et al., Shared molecular neuropathology across major psychiatric disorders parallels polygenic overlap. Science. 359, 693–697 (2018).

6. I. Voineagu et al., Transcriptomic analysis of autistic brain reveals convergent molecular pathology. Nature. 474, 380–384 (2011).

7. N. N. Parikshak et al., Genome-wide changes in lncRNA, splicing, and regional gene expression patterns in autism. Nature. 540, 423–427 (2016).

8. M. J. Gandal et al., Transcriptome-wide isoform-level dysregulation in ASD, schizophrenia, and bipolar disorder. Science. 362, 8127 (2018).

9. G. Ramaswami et al., Integrative genomics identifies a convergent molecular subtype that links epigenomic with transcriptomic differences in autism. Nature Communications. 147, 321–50 (2019).

10. Y. E. Wu, N. N. Parikshak, T. G. Belgard, D. H. Geschwind, Genome-wide, integrative analysis implicates microRNA dysregulation in autism spectrum disorder. Nature Neuroscience. 19, 1463–1476 (2016).

11. W. Sun et al., Histone Acetylome-wide Association Study of Autism Spectrum Disorder. Cell. 167, 1385–1397 (2016).

12. S. Gupta et al., Transcriptome analysis reveals dysregulation of innate immune response genes and neuronal activity-dependent genes in autism. Nature Communications. 5, 1–8 (2014).

13. D. Velmeshev et al., Single-cell genomics identifies cell type–specific molecular changes in autism. Science. 364, 685–689 (2019).

14. M. J. Gandal et al., Broad transcriptomic dysregulation occurs across the cerebral cortex in ASD. Nature. 611, 532–539 (2022).

15. B. B. Lake et al., Neuronal subtypes and diversity revealed by single-nucleus RNA sequencing of the human brain. Science. 352, 1586–1590 (2016).

16. B. B. Lake et al., Integrative single-cell analysis of transcriptional and epigenetic states in the human adult brain. Nature Biotechnology, 1–18 (2017).

17. R. D. Hodge et al., Conserved cell types with divergent features in human versus mouse cortex. Nature. 573, 61–68 (2019).

18. BRAIN Initiative Cell Census Network (BICCN), A multimodal cell census and atlas of the mammalian primary motor cortex. Nature. 598, 86–102 (2021).

19. T. E. Bakken et al., Evolution of cellular diversity in primary motor cortex of human, marmoset monkey, and mouse. bioRxiv, 2020.03.31.016972 (2020).

20. Z. Zhao, A. R. Nelson, C. Betsholtz, B. V. Zlokovic, Establishment and Dysfunction of the Blood-Brain Barrier. Cell. 163, 1064–1078 (2015).

21. C. Escartin et al., Reactive astrocyte nomenclature, definitions, and future directions. Nature Neuroscience. 24, 312–325 (2021).

22. N. A. Oberheim, X. Wang, S. Goldman, M. Nedergaard, Astrocytic complexity distinguishes the human brain. Trends Neurosci. 29, 547–553 (2006).

23. S. Marques et al., Oligodendrocyte heterogeneity in the mouse juvenile and adult central nervous system. Science. 352, 1326–1329 (2016).

24. L. Xiao et al., Rapid production of new oligodendrocytes is required in the earliest stages of motor-skill learning. Nature Neuroscience. 19, 1210–1217 (2016).

25. M. K. Fard et al., BCAS1 expression defines a population of early myelinating oligodendrocytes in multiple sclerosis lesions. Sci. Transl. Med. 9 (2017)

26. A. M. Jurga, M. Paleczna, K. Z. Kuter, Overview of General and Discriminating Markers of Differential Microglia Phenotypes. Front Cell Neurosci. 14, 198 (2020).

27. S. Lodato, P. Arlotta, Generating neuronal diversity in the mammalian cerebral cortex. Annu. Rev. Cell Dev. Biol. 31, 699–720 (2015).

28. B. J. Molyneaux et al., DeCoN: genome-wide analysis of in vivo transcriptional dynamics during pyramidal neuron fate selection in neocortex. Neuron. 85, 275–288 (2015).

29. L. Lim, D. Mi, A. Llorca, O. Marín, Development and Functional Diversification of Cortical Interneurons. Neuron. 100, 294–313 (2018).

30. H. Markram et al., Interneurons of the neocortical inhibitory system. Nat Rev Neurosci. 5, 793–807 (2004).

31. R. Tremblay, S. Lee, B. Rudy, GABAergic Interneurons in the Neocortex: From Cellular Properties to Circuits. Neuron. 91, 260–292 (2016).

32. B. Wamsley, G. Fishell, Genetic and activity-dependent mechanisms underlying interneuron diversity. Nat. Rev. Neurosci. 18, 299–309 (2017).

33. T. Ma et al., Subcortical origins of human and monkey neocortical interneurons. Nature Neuroscience. 16, 1588–1597 (2013).

34. J. T. Morgan et al., Microglial activation and increased microglial density observed in the dorsolateral prefrontal cortex in autism. Biological Psychiatry. 68, 368–376 (2010).

35. D. L. Vargas, C. Nascimbene, C. Krishnan, A. W. Zimmerman, C. A. Pardo, Neuroglial activation and neuroinflammation in the brain of patients with autism. Ann Neurol. 57, 67–81 (2005).

36. S. N. Haber, H. Liu, J. Seidlitz, E. Bullmore, Prefrontal connectomics: from anatomy to human imaging. Neuropsychopharmacology. 47, 20–40 (2022).

37. P. Le Merre, S. Ährlund-Richter, M. Carlén, The mouse prefrontal cortex: Unity in diversity. Neuron. 109, 1925–1944 (2021).

38. A. Kepecs, G. Fishell, Interneuron cell types are fit to function. Nature. 505, 318–326 (2014).

39. Schizophrenia Working Group of the Psychiatric Genomics Consortium, Biological insights from 108 schizophrenia-associated genetic loci. Nature. 511, 421–427 (2014).

40. B. Wamsley et al., Rbfox1 Mediates Cell-type-Specific Splicing in Cortical Interneurons. Neuron. 100, 846–859 (2018).

41. C. K. Vuong et al., Rbfox1 Regulates Synaptic Transmission through the Inhibitory Neuron-Specific vSNARE Vamp1. Neuron. 98, 127–141 (2018).

42. J. Luo, TGF-β as a Key Modulator of Astrocyte Reactivity: Disease Relevance and Therapeutic Implications. Biomedicines. 10, 1206 (2022).

43. E. Lin, P.-H. Kuo, Y.-L. Liu, A. C. Yang, S.-J. Tsai, Transforming growth factor-β signaling pathway-associated genes SMAD2 and TGFBR2 are implicated in metabolic syndrome in a Taiwanese population. Sci Rep. 7, 13589–8 (2017).

44. X. Shen, Y. Qiu, A. E. Wight, H. K. P. O., Definition of a mouse microglial subset that regulates neuronal development and proinflammatory responses in the brain. *Proceedings of the National Academy of Sciences*, PNAS. 119, 21162–41119 (2022).

45. A. Yim, C. Smith, A. M. Brown, Osteopontin/secreted phosphoprotein-1 harnesses glial-, immune-, and neuronal cell ligand-receptor interactions to sense and regulate acute and chronic neuroinflammation. Immunol Rev. 311, 224–233 (2022).

46. S. Aibar et al., SCENIC: single-cell regulatory network inference and clustering. Nat Meth. 14, 1083–1086 (2017).

47. B. Van de Sande et al., A scalable SCENIC workflow for single-cell gene regulatory network analysis. Nat Protoc. 15, 2247–2276 (2020).

48. T. T. Logan, S. Villapol, A. J. Symes, TGF-β superfamily gene expression and induction of the Runx1 transcription factor in adult neurogenic regions after brain injury. PLoS ONE. 8, 59250– (2013).

49. K. M. Dhandapani et al., Astrocyte protection of neurons: role of transforming growth factor-beta signaling via a c-Jun-AP-1 protective pathway. J Biol Chem. 278, 43329–43339 (2003).

50. D. Polioudakis et al., A Single-Cell Transcriptomic Atlas of Human Neocortical Development during Mid-gestation. Neuron. 103, 785–801 (2019).

51. K. Davie et al., A Single-Cell Transcriptome Atlas of the Aging Drosophila Brain. Cell. 174, 982–998 (2018).

52. X. Han et al., Construction of a human cell landscape at single-cell level. Nature. 581, 303–309 (2020).

53. M. Bentsen et al., ATAC-seq footprinting unravels kinetics of transcription factor binding during zygotic genome activation. Nature Communications. 11, 4267–11 (2020).

54. F. Tian et al., Core transcription programs controlling injury-induced neurodegeneration of retinal ganglion cells. Neuron. 110, 2607–2624.e8 (2022).

55. J. Grove et al., Common risk variants identified in autism spectrum disorder, Nat Genet. 51, 431–444 (2019).

56. F. K. Satterstrom et al., Large-Scale Exome Sequencing Study Implicates Both Developmental and Functional Changes in the Neurobiology of Autism. Cell. 180, 568–584 (2020).

57. C. S. Leblond et al., Operative list of genes associated with autism and neurodevelopmental disorders based on database review. Mol Cell Neurosci. 113, 103–623 (2021).

58. J. C. Darnell et al., FMRP Stalls Ribosomal Translocation on mRNAs Linked to Synaptic Function and Autism. Cell. 146, 247–261 (2011).

59. C. Li, J. S. Fleck, C. Martins-Costa, T. B., Single-cell brain organoid screening identifies developmental defects in autism. Biorxiv, 2022.09.15.508118 (2022).

60. A. Gordon et al., Long-term maturation of human cortical organoids matches key early postnatal transitions. Nature Neuroscience. 24, 331–342 (2021).

61. X. Jin et al., In vivo Perturb-Seq reveals neuronal and glial abnormalities associated with autism risk genes. Science. 370 (2020).

62. R. L. Walker et al., Genetic Control of Expression and Splicing in Developing Human Brain Informs Disease Mechanisms. Cell. 179, 750–771 (2019).

63. S. Khan et al., Local and long-range functional connectivity is reduced in concert in autism spectrum disorders. *Proceedings of the National Academy of Sciences*, PNAS. 110, 3107–3112 (2013).

64. M. Kikuchi et al., Reduced long-range functional connectivity in young children with autism spectrum disorder. Soc Cogn Affect Neurosci. 10, 248–254 (2015).

65. D. H. Geschwind, P. Levitt, Autism spectrum disorders: developmental disconnection syndromes. Curr. Opin. Neurobiol. 17, 103–111 (2007).

66. S. J. Sanders et al., De novo mutations revealed by whole-exome sequencing are strongly associated with autism. Nature. 485, 237–241 (2012).

67. I. Iossifov et al., The contribution of de novo coding mutations to autism spectrum disorder. Nat Genet. 515, 216–221 (2014).

68. Christensen D, Zubler J., From the CDC: Understanding Autism Spectrum Disorder. Am J Nurs. 10, 30–37 (2020).

69. L. de la Torre Ubieta, H. Won, J. L. Stein, D. H. Geschwind, Advancing the understanding of autism disease mechanisms through genetics. Nat Med. 22, 345–361 (2016).

70. D. M.-C. L. PhD, M. V. L. PhD, P. S. B.-C. PhD, Autism. The Lancet. 383, 896–910 (2014).

71. T. Stuart et al., Comprehensive Integration of Single-Cell Data. Cell. 177, 1888–1902 (2019).

72. C. W. Law, Y. Chen, W. Shi, G. K. Smyth, voom: Precision weights unlock linear model analysis tools for RNA-seq read counts. Genome Biol. 15, R29–17 (2014).

73. E. J. Rossin et al., Proteins Encoded in Genomic Regions Associated with Immune-Mediated Disease Physically Interact and Suggest Underlying Biology. PLoS Genet. 7, 1001273–13 (2011).

74. H. K. Finucane et al., Partitioning heritability by functional annotation using genome-wide association summary statistics. Nat Genet. 47, 1228–1235 (2015).

75. T. Stuart, A. Srivastava, S. Madad, C. A. Lareau, R. Satija, Single-cell chromatin state analysis with Signac. Nat Meth. 18, 1333–1341 (2021).

